# Heme-stress activated NRF2 signaling skews fate trajectories of bone marrow cells from dendritic cells towards red pulp-like macrophages

**DOI:** 10.1101/2021.07.29.454342

**Authors:** Florence Vallelian, Raphael M. Buzzi, Marc Pfefferlé, Ayla Yalamanoglu, Andreas Wassmer, Thomas Gentinetta, Kerstin Hansen, Rok Humar, Nadja Schulthess, Dominik J. Schaer

**Affiliations:** Division of Internal Medicine, University of Zurich, Switzerland; CSL Behring AG, Research and Development, Bern, Switzerland

## Abstract

Heme is an erythrocyte-derived toxin that drives disease progression in hemolytic anemias, such as sickle cell disease. During hemolysis, specialized bone marrow-derived macrophages with a high heme-metabolism capacity orchestrate disease adaptation by removing damaged erythrocytes and heme-protein complexes from the blood and supporting iron recycling for erythropoiesis. Since chronic heme-stress is noxious for macrophages, erythrophagocytes in the spleen are continuously replenished from bone marrow-derived progenitors. Here, we hypothesized that adaptation to heme stress progressively shifts differentiation trajectories of BM progenitors to expand the capacity of heme-handling monocyte-derived macrophages at the expense of the homeostatic generation of dendritic cells, which emerge from shared myeloid precursors. This heme-induced redirection of differentiation trajectories may contribute to hemolysis-induced secondary immunodeficiency. We performed single-cell RNA sequencing with directional RNA velocity analysis of GM-CSF-supplemented mouse bone marrow cultures to assess myeloid differentiation under heme stress. We found that heme-activated NRF2 signaling shifted the differentiation of bone marrow cells towards antioxidant, iron-recycling macrophages, suppressing the generation of dendritic cells in heme-exposed bone marrow cultures. Heme eliminated the capacity of GM-CSF-supplemented bone marrow cultures to activate antigen-specific CD4 T cells. The generation of functionally competent dendritic cells was restored by NRF2 loss. The heme-induced phenotype of macrophage expansion with concurrent dendritic cell depletion was reproduced in hemolytic mice with sickle cell disease and spherocytosis and associated with reduced dendritic cell functions in the spleen. Our data provide a novel mechanistic underpinning of hemolytic stress as a driver of hyposplenism-related secondary immunodeficiency.

**GRAPHICAL ABSTRACT:** 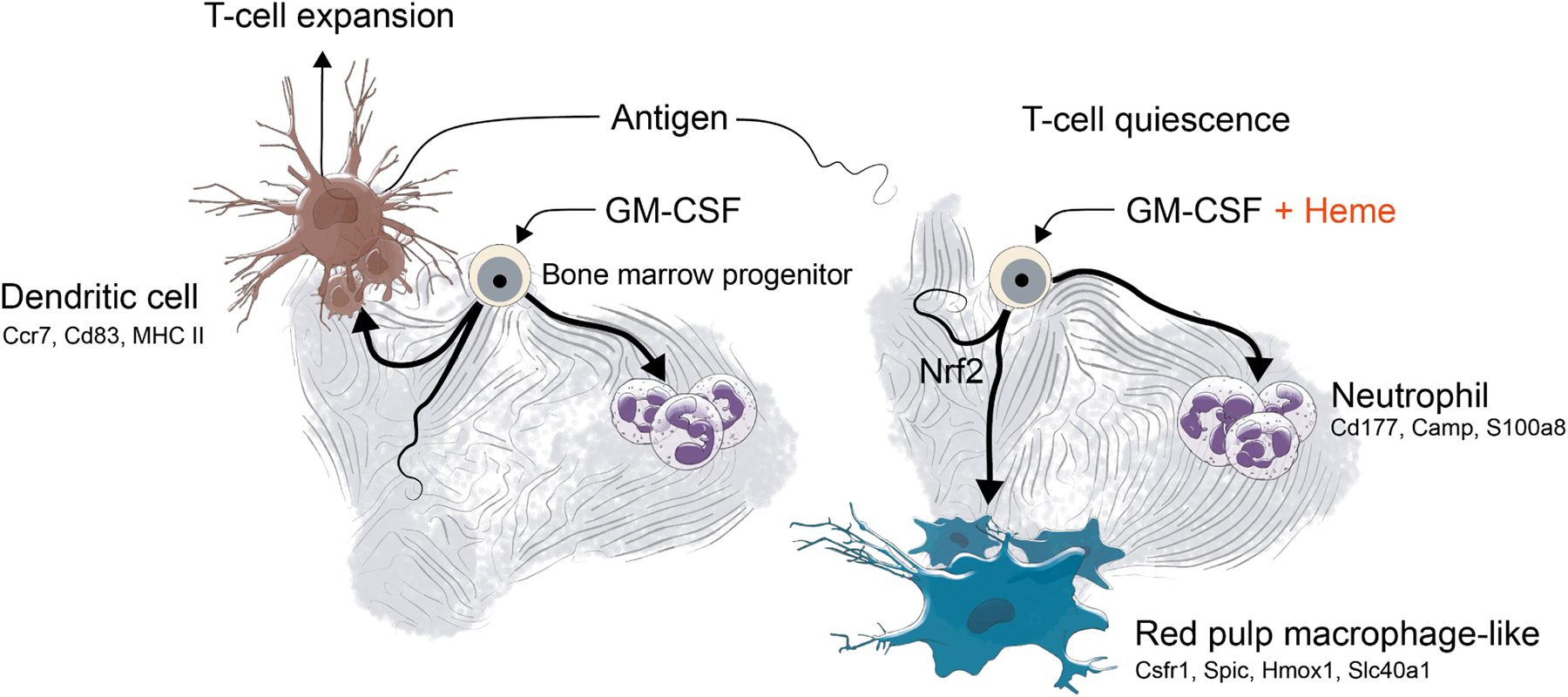

## INTRODUCTION

The hemolytic destruction of red blood cells (RBCs) generates large quantities of cell-free hemoglobin and heme, which accumulate in the blood and tissues. These RBC-derived toxins drive morbidity and mortality in hemolytic anemias via a number of pathophysiological pathways that cause nitric oxide depletion, lipid peroxidation, and heme engagement of cellular signaling receptors ^1–4^. While primary heme toxicities are localized in the cardiovascular system and kidneys, more recent research has identified heme as a context-dependent stimulator and suppressor of immunity and inflammation ^3–5^. Immunosuppressive functions of heme may contribute towards a poorly understood secondary immunodeficiency, called hyposplenism, which is a critical contributor to mortality in patients with genetic hemolytic anemia ^6–9^.

Hemolytic anemias such as sickle cell disease (SCD), thalassemia, and spherocytosis are the most frequent monogenic diseases worldwide and have exerted strong evolutionary pressure on physiological adaptation enhancing disease tolerance to anemia and heme-stress ^10, 11^. A key role of this adaptation is provided by mononuclear phagocytes in the liver and spleen. These macrophages remove damaged RBCs and toxic heme-protein complexes from the blood, which limits hemolysis-related tissue toxicity and ultimately sustains iron metabolism and erythropoiesis ^12–16^. However, chronic heme stress, such as that in hemolytic anemias, is noxious for erythrophagocytic macrophages, and their continuous replacement by bone marrow (BM)-derived monocytes is thus needed to maintain homeostasis ^17–22^. This replacement is driven by a regulatory network that includes the heme-activated transcription factor Spi-C ^21^ and PPARγ ^23^. The antioxidant and iron-recycling functions are further supported by ATF1 ^24^ and the basic leucine zipper (bZIP) protein transcription factor NFE2L2 (NRF2), which is known as the master regulator of the cellular response to oxidative stress ^25^.

Like monocyte-derived macrophages, dendritic cells (DCs) emerge from myeloid-rooted BM progenitors. DCs are the professional antigen-presenting cells and are essential for the initiation of effector T cell responses ^26^. Despite their key role at the interface of innate and adaptive immunity, there is a lack of research defining the influence of hemolysis and heme stress on DCs and how a potential deregulation of DC homeostasis could contribute to hemolysis-associated immunodeficiency. Similar to heme-handling macrophages in the hemolytic spleen, DCs are short-lived cells that continuously emerge from the BM and differentiate into mature antigen-presenting cells in tissues ^26–28^. Monocyte-derived macrophages and DCs are progenies of common myeloid BM-precursors, which implies a potential competition of macrophage and DC generation. ^29–31^. Based on this prior knowledge, we reasoned that adaptation to chronic heme stress may progressively shift the differentiation trajectories of myeloid BM progenitors to expand the capacity of heme-handling macrophages at the expense of homeostatic DC production. Although such a hematopoietic deviation may support adaptive physiology in hemolytic stress, the resulting deficiency in DCs may contribute to hyposplenism.

Therefore, to explore how heme-stress could perturbate macrophage and DC homeostasis, we performed time-resolved single-cell RNA-sequencing (scRNA-seq) experiments with mouse BM cultures. We found that heme redirected the differentiation trajectory of BM cells toward antioxidant and iron-recycling macrophages rather than DCs via an NRF2-dependent signaling pathway. Heme stress in mice with SCD replicated this phenotype in vivo and showed the expansion of heme-handling macrophages and a concomitant reduction in DCs with an impaired capacity to prime antigen specific T cells. Our data reveal a new regulatory function of heme in the immune system and points to a novel mechanistic understanding of secondary immunodeficiency in hemolytic anemia.

## RESULTS

### Heme exposure suppressed the phenotype of functional DCs in mouse BM cultures

To investigate the impact of heme-exposure on myeloid cell differentiation trajectories in vitro, we cultured lineage-depleted (Lin^—^) BM cells from C57BL/6 mice supplemented with recombinant M-CSF or GM-CSF, which are known to induce BM cells into macrophages and DCs, respectively. Within seven days, these M-CSF- and GM-CSF BM cultures generated populations of CD11b^+^/CD115^+^/MHC-class II^low^ macrophages and CD11c^+^/CD115^—^/MHC-class II^high^ DC-like cells, respectively (**Supplementary Figure 1A**) ^29, 32^. We characterized the potential heme-handling function of these cells by measuring the uptake of SnMP, a fluorescent heme-analog, using a spectral flow analyzer. In contrast to the DC-like cells, we detected a strong SnMP-typical fluorescence signal in the macrophages, suggesting functional heterogeneity (**Supplementary Figure 1B**).

To fully characterize the cell type heterogeneity and changes induced by heme under the two culture conditions, we exposed M-CSF BM cultures and GM-CSF BM cultures to heme-albumin (heme) or albumin (vehicle control) and then performed a multiplexed scRNA-seq experiment (**Figure 1A**). Cells under each condition were labeled with DNA-barcoded antibodies, pooled and processed for sequencing. **Figure 1A** shows a uniform manifold approximation and projection (UMAP) of the 4,852 cells merged across the four treatment conditions (growth factor x treatment). The 15 clusters were assigned to cell types using an unbiased approach with an algorithm that performs a per-cluster search for populations with similar gene expression in a database of > 15 million annotated single-cell transcriptomes ^33^ (**Supplementary Figure 1C**). The M-CSF BM cultures generated only macrophages, with minor database hits consistent with monocytes and microglia. In contrast, the GM-CSF BM cultures were more heterogeneous and consisted of macrophages, DCs and neutrophils (**Figure 1B**). Split UMAP plots for the four conditions showed clear segregation of the GM-CSF and M-CSF BM cultures and also of the control and heme-exposed cells, which implies a strong effect of heme-exposure on the overall gene expression profile (**Figure 1C**).

**Figure 1:**
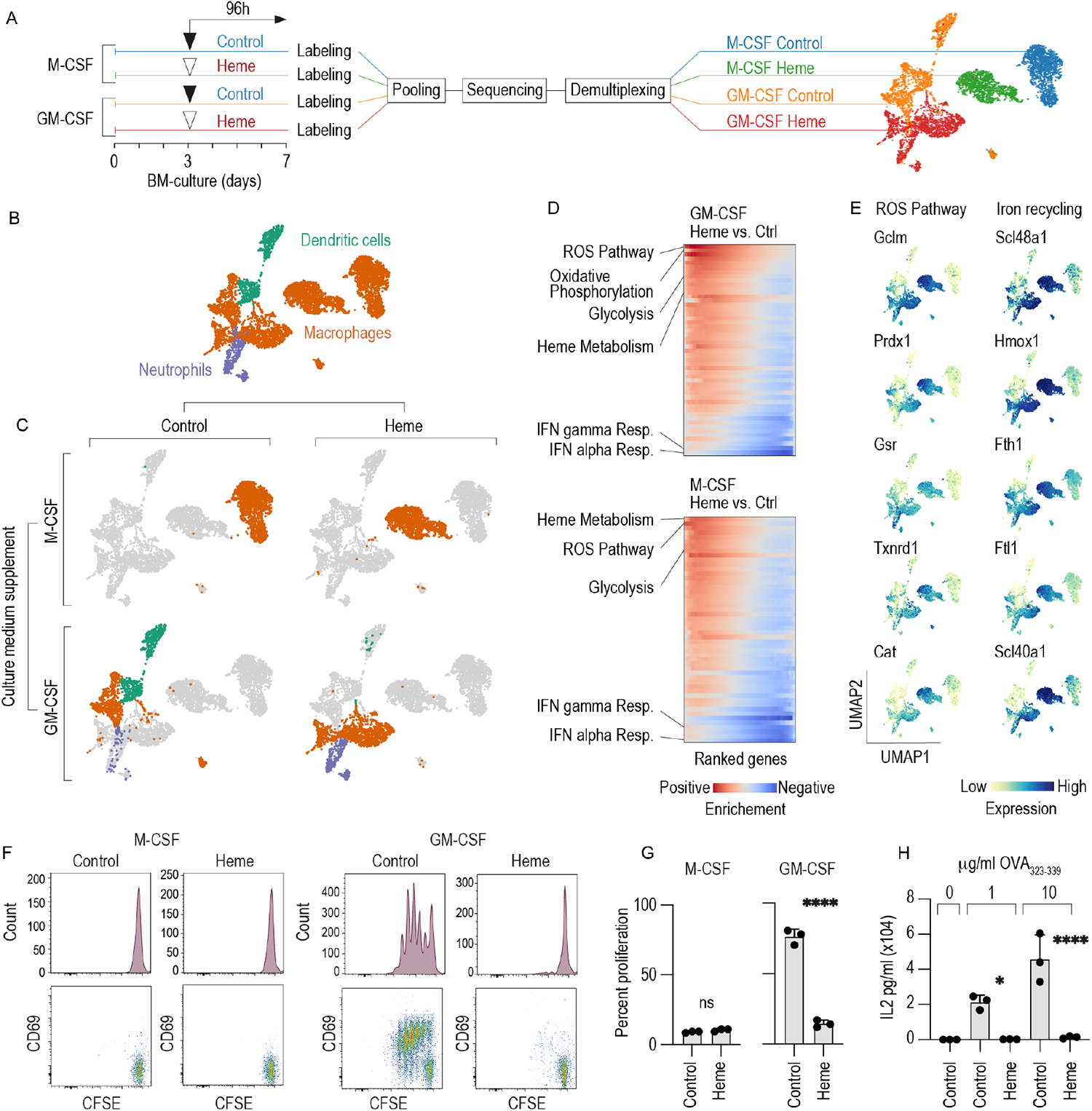
Heme exposure suppressed CD11c^+^/MHC class II^high^/CD115 (Csf1R)^—^ DC phenotype and function in GM-CSF mouse BM cultures. A. Left: Outline of the multiplexed scRNA-seq experiment. M-CSF- and GM-CSF BM cultures were stimulated with heme (300 µM) or vehicle on day 3. On day 7, the cells were harvested, tagged with oligo-barcoded antibodies, and pooled into a single scRNA-seq sample for sequencing. Demultiplexing and reassignment of treatment groups were performed after quality assessment, filtering, and cell-type attribution. Right: UMAP plot showing cells colored by DNA-barcoded antibodies (hashtag labeling). B. UMAP plot showing cells colored by cell type. C. Factorial UMAP plots of the dataset separated by culture supplement (M-CSF versus GM-CSF) and treatment condition (control versus heme exposure). The data points are colored by cell type (orange=macrophages, purple=neutrophils, green=DCs). D. GSEA to identify the differentially expressed genes (heme exposure versus control) in GM-CSF or M-CSF BM cultures. The heatmaps represent the running enrichment score per gene set category (red for positive enrichment, blue for negative enrichment). Each row represents one gene set category. E. UMAP plots showing the expression levels of the top five enriched genes defining the ROS pathway and genes associated with iron recycling in the whole data set. F. M-CSF- and GM-CSF BM cells were cultured with heme (300 µM) or vehicle. On day 7, the cells were harvested, pulsed with OVA_323-339_ (1 µg/ml), washed to remove any remaining heme, and then incubated with CFSE-labeled naïve CD4^+^ T cells isolated from OT-2 mice in a 1/5 BM cells to T cells ratio. The proliferation of CD4^+^ T cells was assessed by evaluating the degree of CFSE dilution measured by flow cytometry after three days in coculture. The cells were also stained for the T cell activation marker CD69. Representative data for cells gated based on positive CD4 expression are shown. G. Cumulative T cell proliferation data from three independent coculture experiments. H. GM-CSF BM cells were cultured with heme (300 µM) or vehicle. On day 7, the cells were harvested, pulsed with OVA_323-339_ (1 or 10 µg/ml), washed to remove any remaining heme, and then incubated with CFSE-labeled naïve CD4^+^ T cells. The proliferation of CD4^+^ T cells was assessed after three days in coculture by flow cytometry. Cumulative T cell proliferation data from three independent coculture experiments are shown. The data in G and H are presented as the means ± SDs. Each dot represents one independent experiment. t-test (G); one-way ANOVA with Tukey’s multiple comparison test (H); n.s. = not significant, \**P* ≤ 0.05, \*\**P* ≤ 0.01, \*\*\**P* ≤ 0.001, \*\*\*\**P* ≤ 0.0001.

The key strength of this experiment was the precise definition of how the presence of heme changed the cell types that were generated. In the M-CSF BM cultures all cells were assigned a macrophage phenotype regardless of whether heme was present in the culture or not. In contrast and unexpectedly, in the GM-CSF BM cultures, we found that the prevailing DC population represented by clusters 12 and 13 completely disappeared in the presence of heme, whereas a small population exhibiting neutrophil-specific gene expression emerged in only the heme-treated cells (cluster 14) (**Figure 1C, Supplementary Figure 1C**). The expression of canonical marker genes for DCs ^34, 35^, macrophages, and neutrophils is shown in **Supplementary Figure 1D**.

To identify functional implications of the overall gene expression changes induced by heme, we performed gene set enrichment analysis (GSEA) of differentially expressed genes identified from a pseudo-bulk analysis of heme-treated versus control samples (**Figure 1D** and **Supplementary Figure 2A**). In both GM-CSF- and M-CSF BM cultures, heme strongly induced the hallmark gene set for the reactive oxygen species (ROS) pathway including the antioxidant genes Gclm, Prdx1, Gsr, Txnrd1, and Cat (**Figure 1E**). We also found that heme enhanced expression of the iron-recycling machinery, defined by the heme transporter Slc48a1, the heme-degradation enzyme heme oxygenase (Hmox1), and genes related to iron storage (Ftl1, Fth1) and iron export (Slc40a1) (**Figure 1E**). In contrast, we found consistent suppression (i.e., negative enrichment) of the IFN alpha and IFN gamma responses in the heme-treated cells compared to the control cells (**Figure 1D** and **Supplementary Figure 2A**). Collectively, these GSEA results suggest that heme enhances antioxidant and iron-recycling functions, while specific immune functions may be suppressed.

The defining immune function of DCs is their capacity for antigen-specific priming of naïve T cells ^26, 36^. Therefore, to functionally validate our scRNA-seq study, we analyzed the proliferation and activation of CD4^+^ T cells isolated from OVA-specific recombinant T cell receptor transgenic mice (OT-2 mice) in cocultures with ovalbumin (OVA_323-339_) peptide-pulsed BM cells by evaluating carboxyfluorescein succinimidyl ester (CFSE) dilution and cytokine production, respectively. We performed these experiments under the assumption that the observed T cell response would directly reflect the output of mature and functionally competent DCs in the different cell cultures. Importantly, before the coculture with T cells was initiated, the BM cultures were washed to avoid direct interference of heme with T cell activation and proliferation. Consistent with our phenotype-based definition derived from the scRNA-seq data (**Figure 1C**), coculture of T cells with GM-CSF BM cultures, but not M-CSF BM cultures resulted in the appearance of multiple CD69^+^T cell populations with progressively reduced (i.e., diluted) CFSE fluorescence, indicating activation and division of OT-2 CD4^+^ T cells (**Figure 1F** and **G**). This activation was accompanied by the secretion of IL-2, which was dose-dependent in respect to the OVA_323-339_ peptide concentration used to pulse BM cultures (**Figure 1H**). Heme exposure completely suppressed the CD4^+^ T cell-priming capacity of GM-CSF BM cultures (**Figure F, G, H**), which was consistent with the heme-suppressed DC phenotype (**Figure 1D**). To verify that this effect of heme is maintained in vivo, we adoptively transferred CFSE-labeled OT-2 CD4^+^ T cells and OVA_323-339_ peptide-pulsed GM-CSF BM cells that were cultured with heme or vehicle into CD45.1 Rag1^−/−^ mice and analyzed the proliferation of the T cells in the spleen after three days by flow cytometry. In this experiment, DC retained a strong antigen presentation capacity that was profoundly suppressed by heme (**Supplementary Figure 1E and F**). In summary, these experiments indicated that heme-stress suppressed the capacity of GM-CSF BM cultures to induce activation and proliferation of antigen-specific T cells consistent with the disappearance of DCs.

Collectively, based on these experimental results, we hypothesized that heme exposure redirects BM cell differentiation trajectories in favor of antioxidant and iron-recycling macrophages at the expense of functionally mature DCs. Furthermore, we found that cell-type heterogeneity in the GM-CSF BM cultures provides a unique opportunity to explore heme-deviated myeloid cell differentiation in a single culture dish, which prompted the use of only GM-CSF BM cultures in the remainder of the study.

### Heme skewed the cell differentiation fates of mouse BM cultures from DCs towards macrophages

To capture the cell differentiation trajectories of GM CSF BM cultures and the effect of heme in more detail, we extended our studies with another scRNA-seq experiment using a time-resolved scRNA-seq experiment, as outlined in **Figure 2A**. Briefly, we cultured Lin^—^ BM cells with GM-CSF, added heme or vehicle on day 3 of culture, and cultured the cells for an additional 24, 72, or 96 h. Using CellRank ^37^, we combined transcriptional similarity data with directional RNA velocity information to delineate three terminal differentiation states, or macrostates (**Figure 2B**). CellSearch similarity matching attributed these macrostates to DCs, macrophages, and neutrophils (**Figure 2C**), which is consistent with the expression of canonical marker genes für these cell types shown in **Figure 2D**. Additionally, we identified three transitional cell states consistent with progenitors and cells at an intermediate differentiation state (**Figure 2E**). **Figure 2F** shows the UMAP plots obtained in the experiment with data from cells under each treatment condition for each exposure time viewed separately. Over time, the transcriptome profiles progressed along the RNA velocity vectors calculated for the whole dataset (**Figure 2E**), with the heme-treated and control cultures showing very different outcomes. Cells in the control culture initially progressed on day 3 along two trajectories towards neutrophils and DCs. At the end of the study, on day 7, the neutrophil arm had disappeared, leaving a prevailing population of DCs (brown) alongside an intermediate state population with marker gene expression consistent with immature DCs and macrophages/monocytes. In the heme-exposed BM culture, the propagation and collapse of the neutrophils were comparable to those in the control culture, but the final differentiation fate was strongly skewed away from DCs towards Spic^+^ red pulp-like macrophages, with high expression of key heme and iron metabolism genes such as the heme transporter Slc48a1, Hmox1, ferritin genes (Fth1, Flt1), and the iron exporter Slc40a1 (**Supplementary Figure 2B**). This qualitative interpretation was supported by a quantitative analysis of the discrete cell state proportions (**Figure 2G**) and by the day-wise cumulative absorption probabilities (**Figure 2H**), which represented the probabilistic fate that any cell within the experiment would differentiate into one of the three terminal states representing neutrophils, macrophages, or DCs. Here, the highest absorption probability on day 7 suggested a preferential DC cell fate for cells in the control condition and a preferential macrophage cell fate for cells in the heme-treated condition. Individual cell absorption probabilities for all cells across timepoints and treatments are shown in **Figure 2I** ^37, 38^.

**Figure 2:**
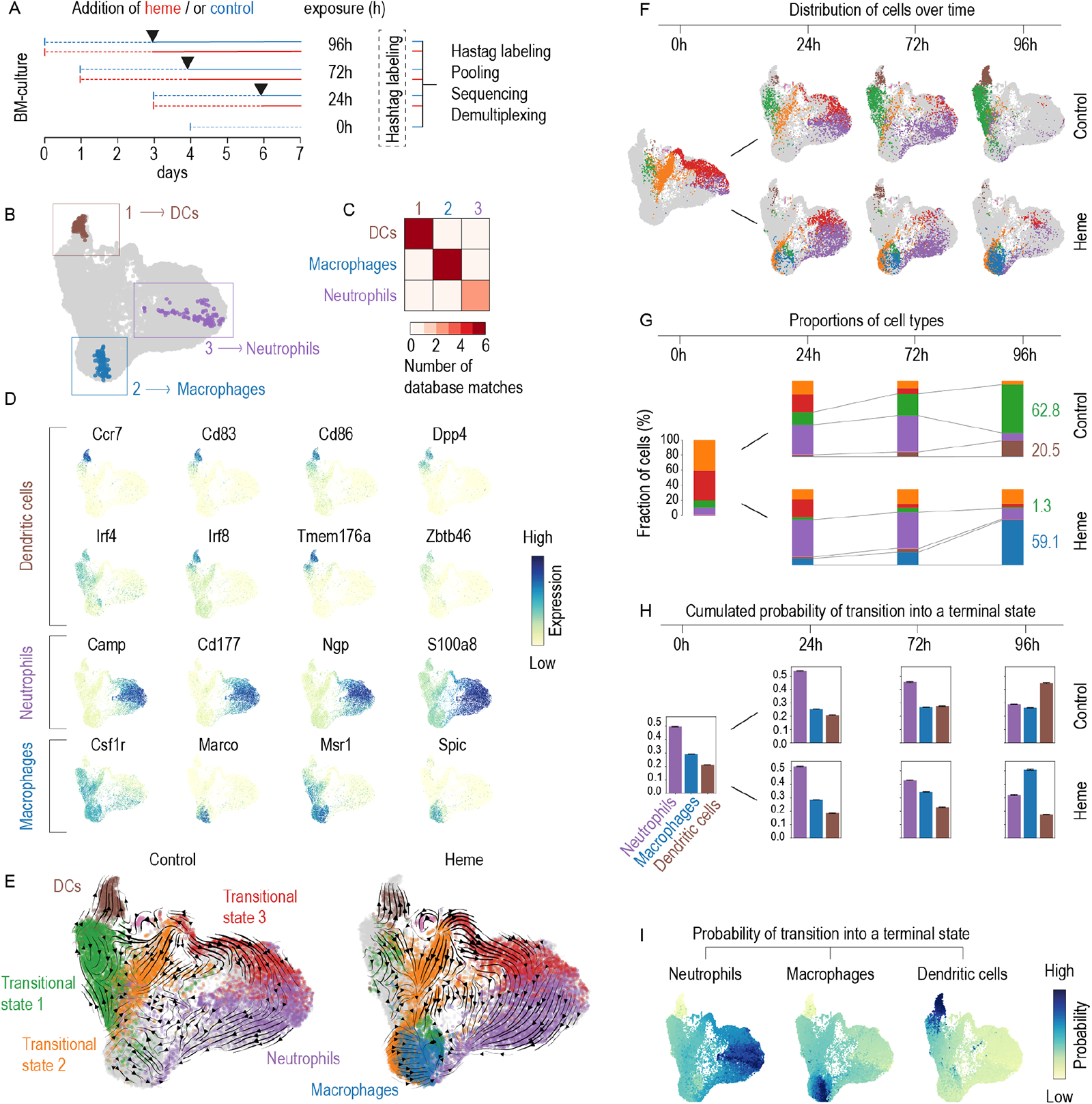
Heme redirects the cell differentiation pathways in GM-CS mouse BM cultures from DCs towards macrophages. A. Outline of the multiplexed scRNA-seq experiment. GM-CSF BM-cultures were initiated on day 0, 1, 3 and 4. Each culture was treated with heme (300 µM) or vehicle control (black arrows) on day 3 and cells were harvested on day 7 providing heme exposure periods of 0, 24, 72, and 96 h. After, the cells were harvested, tagged with DNA-barcoded antibodies, pooled, and processed for sequencing. After quality assessment, filtering, and cell type attribution, the experiment was demultiplexed to reassign individual cells to their sample of origin. B. UMAP plot of the scRNA-seq results highlighting the cells that were automatically selected by the CellRank algorithm as most representative of one of three, yet undefined, terminal differentiation states (1 brown, 2 blue, 3 violet). C. The terminal state-typical cell populations selected in (B) were used in a similarity search against > 15 million annotated cells from published studies. Similarity matching was performed using CellSearch (Bioturing). For each macrostate (1, 2 or 3), the heatmap shows the number of high-quality matches for DCs, macrophages, and neutrophils that were then used for cell-type attribution. D. Expression intensity projections of the scRNA-seq data showing canonical marker genes for cDCs, neutrophils, and macrophages. E. UMAP plots of the scRNA-seq data separated by treatment status (control and heme). The superimposed arrows indicate RNA velocity vectors that were calculated for each treatment condition across time points. The colors represent the three terminal states and three additional transition states. The macrostate colors will be used for the remainder of the figure. F. UMAP plots of the scRNA-seq data separated by heme treatment status and time point illustrate the treatment-dependent differentiation pathways across the four time points. G. Stack bar charts illustrating discrete cell-type attributions as a proportion of the whole number of cells per sample,for each time and treatment condition. Numbers are percentages. H. Cumulative absorption probabilities summarizing the probability of each cell reaching each terminal state defined as DCs, macrophages, and neutrophils given for each time point for both experimental conditions I. UMAP plots with the cells colored according to their probability of reaching the terminal state defined as DCs, macrophages, or neutrophils. Data of all treatment conditions (heme x time) are merged.

Collectively, these results reveal that heme exposure skewed the probabilistic differentiation fates of GM-CSF BM cultures from DCs and towards macrophages with a red pulp macrophage-like gene expression profile to support heme metabolism and iron recycling.

### Suppression of DCs in mouse BM cultures due to heme exposure proceeds via the transcription factor NRF2

Next, we aimed to identify the signaling pathway through which heme suppresses the generation of mature DCs. We performed a transcription factor-binding site enrichment analysis with the differential gene expression data from the heme-exposed versus control GM-CSF BM cultures exposed in Figure 1 using EnrichR ^39^ and the TRRUST gene-set library ^40^. NFE2L2 (NRF2) was found to be the most significant hit (**Figure 3A**).

**Figure 3:**
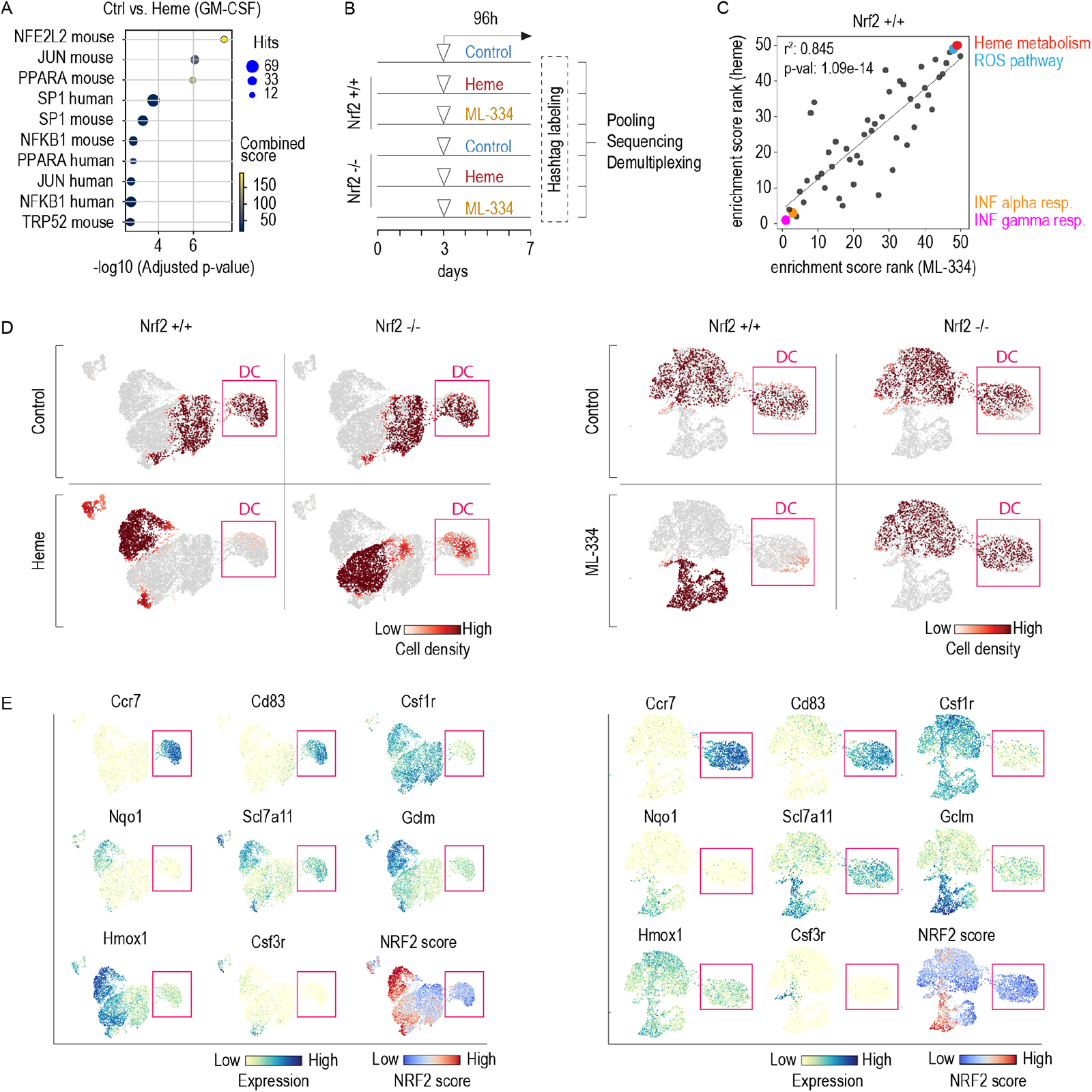
Heme-NRF2 signaling impaired DC differentiation in GM-CSF mouse BM cultures. A. Dot plot showing the top 10 enriched terms from an EnrichR analysis (using the TRRUST gene-set library) of the genes upregulated by heme versus vehicle treatment (log2FC > 1, *p*-value < 0.05) in the GM-CSF BM culture experiment shown in Figure 1. The size of the dots represents the number of overlapping genes, and the color represents the combined score. The enriched terms in the columns of the matrix are ranked based on their adjusted *p*-value. B. scRNA-seq analysis of GM-CSF BM cultures performed with BM from Nrf2^+/+^ and Nrf2^-/-^ mice. The cells were treated on day 3 with vehicle, heme (300 µM) or ML-334 (10 µM). On day 7, each sample was tagged with DNA-barcoded antibodies and pooled for sequencing and downstream analysis. C. Correlation plot of the ranked GSEA enrichment scores for heme-regulated and ML-334-regulated gene sets. GSEA was performed using differential gene expression data from scRNA-seq analysis of Nrf2^+/+^GM-CSF BM cultures treated with heme or ML-334 versus control. D. Factorial scRNA-seq analysis of GM-CSF BM cultures performed with BM from Nrf2^+/+^ and Nrf2^-/-^ mice. The UMAP plots are colored to represent the cell densities (red) for each experimental condition (left panel: control, heme; right panel: control, ML-334). Pink box highlights DC population. This illustration indicates that heme and ML-334 suppress the generation of DCs in Nrf2^+/+^ but not Nrf2^-/-^ BM cultures. E. Expression intensity projections of selected marker genes in combined Nrf2^+/+^ and Nrf2^-/-^ BM cells treated with vehicle/heme (left) or vehicle/ML-334 (right). DCs were identified as the Ccr7^high^, Cd83^high^, and Csf1r**^—^** cell population (highlighted by pink box). The UMAP on the lower right displays the scored expression of the full Nrf2-regulated gene set from the TRRUST gene-set library. A high NRF2 score was detected in Nrf2^+/+^ cells treated with heme or ML-334.

To validate the function of NRF2, we performed a two-factorial multiplexed scRNA-seq experiment using GM-CSF BM cultures from Nrf2*^-/-^* mice and wild-type littermates that were treated with heme or with the selective NRF2 activator ML-334 (**Figure 3B**). As an initial screen, we compared differential gene expression data of the compound-treated (heme or ML-334) versus vehicle-treated wild-type cells after functional consolidation of the data by GSEA. **Figure 3C** demonstrates the high correlation of the ranked GSEA results for heme and ML-334, respectively. Under both treatment conditions, genes involved in heme metabolism and the ROS pathway were strongly upregulated, while those involved in IFN signaling pathways were suppressed to the greatest degree, suggesting that selective NRF2 activation replicates the effect of heme on GM-CSF BM cultures.

**Figure 3D** shows UMAP plots of the scRNA-seq experiment for the cell densities under each experimental condition separated by genotype and treatment. Expression heatmaps of canonical marker genes of mature DCs (Ccr7, Cd83), monocytes-macrophages (Csf1r), and neutrophils (Csf3r) and that of representative genes known to be regulated by NRF2 (Gclm, Nqo1, Scl7a11, and Hmox1) are highlighted in **Figure 3E**, and these results are accompanied by a cumulative NRF2 target gene expression score, which compares the mean expression of the Nrf2 target genes from the TRRUST gene-set library to a randomly selected reference set. The data indicated that both heme and ML-334 activated Nrf2-related gene expression in wild-type cells and almost completely blocked the emergence of Ccr7^+^ Cd83^+^ Csf1r^—^ DCs. Nrf2 deficiency rescued the generation of DCs in heme-treated and ML-334-treated cells, which suggests that a linked heme-NRF2 signaling pathway suppressed DC generation in the heme-exposed BM cultures.

Collectively, the scRNA-seq results indicate that heme-activated NRF2 suppresses the emergence of mature DCs from BM progenitors.

### Cell-intrinsic heme-NRF2 signaling suppressed DC and functional antigen-presentation capacity in mouse BM cultures

In the next step, we validated the function of the heme-NRF2 signaling pathway as a suppressor of DC generation by flow cytometry and T cell proliferation assays. For these studies we treated GM-CSF BM cultures with different concentrations of heme and selective NRF2 activators for four days. The results demonstrated that the absence of the Nrf2 gene blocked the heme-induced loss of CD11c^+^/MHC class II^high^/CD115 (Csf1R)^—^ DCs (**Figure 4A** and **B**). Additionally, the two NRF2 activators ML-334 and RA-839 did not suppress differentiation into the DCs in Nrf2^—/—^ cells (**Figure 4C** and **D**), confirming the target-specificity of the observed effect. We also performed heme treatment experiments with mixed cultures of Nrf2^—/—^ and wild-type BM cells in the same cell culture dish, exploiting the congenic CD45.1/CD45.2 system, which can be used with flow cytometry to discriminate cell origin ^41^ (**Figure 4E**). The results showed that in a mixed culture heme selectively suppressed the appearance of wild-type (CD45.1^+^) but not Nrf2^—/—^ (CD45.2^+^) CD11c^+^/CD115 (Csf1R)^—^ DCs. These experiments suggested that heme-NRF2 signaling functions as a cell-intrinsic pathway and exclude significant paracrine effects of NRF2-regulated metabolites or cytokines (**Figure 4F**). Nrf2 gene knockout also completely restored the impaired antigen-presentation capacity of heme- and ML-334-treated BM cultures (**Figure 4G** and **H**).

**Figure 4:**
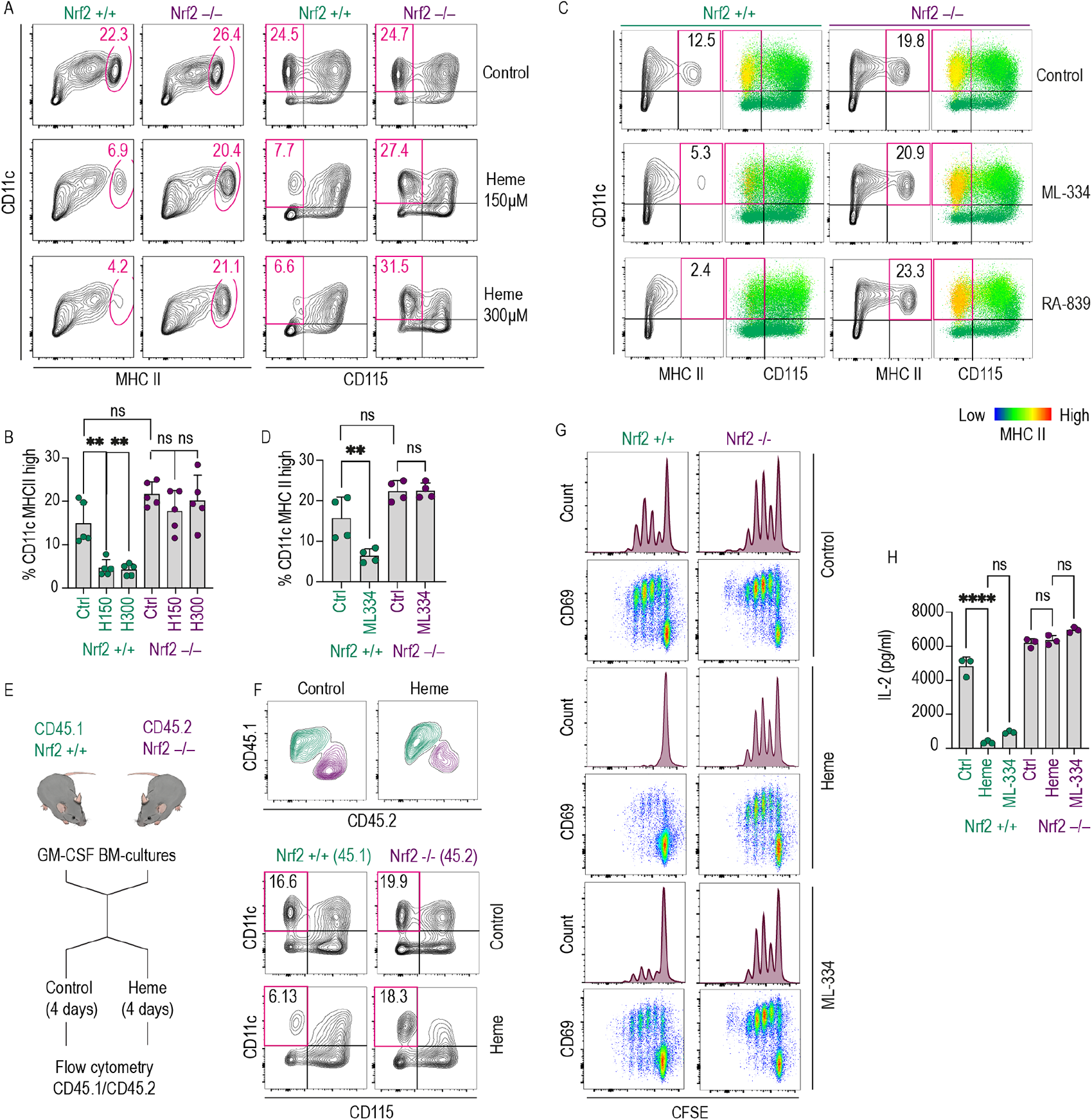
The emergence of functional DCs was suppressed by cell-intrinsic heme-NRF2 signaling in GM-CSF mouse BM cultures. A. Flow cytometry contour plots of Nrf2^+/+^ and Nrf2^-/-^ GM-CSF BM cultures treated with heme (150 or 300 µM) or vehicle for 96 h (culture days 3-7). Based on the scRNA-seq data shown in Figure 1, we defined DCs in this culture as CD11c^+^/MHC class II^high^ or CD11c^+^/CD115^—^ (boxed in red). B. Proportion of DCs defined as CD11c^+^ MHC class II^high^ across treatment conditions and genotypes. C. Flow cytometry contour plots of Nrf2^+/+^ and Nrf2^-/-^ GM-CSF BM cultures treated with the Nrf2 activator ML-334 (10 µM) or RA-839 (15 µM) for 96 h. The data illustrate the Nrf2-dependent depletion of MHC class II^high^ (yellow) cells in the CD11c^+^ CD115^—^ population. D. Proportions of DCs defined a CD11c^+^ MHC class II^high^ across treatment conditions and genotypes. E. Outline of BM coculture experiments: Nrf2^-/-^ (CD45.2) and wild-type (CD45.1) BM cells were cocultured in the same dish and treated with heme-albumin or vehicle for 96 h. During the final analysis, the original genotype of each cell was assigned by flow cytometric analysis of the congenic CD45.1/CD45.2 markers. F. Flow cytometry contour plots of a representative coculture experiment. Heme exposure leads to selective depletion of CD11c^+^ CD115^—^ DCs in the wild-type (CD45.1) cell population but not in the Nrf2-knockout cell population. G. Nrf2^+/+^ and Nrf2^-/-^ GM-CSF BM cultures treated with vehicle, heme-albumin (150 µM), or ML-334 (10 µM) for 96 h were pulsed with OVA_323-339_ (1 µg/ml), washed to remove heme and excess peptide, and cocultured with CFSE-labeled OT-2 CD4^+^ T cells. The dilution of CFSE after four days was measured by flow cytometry. The cells were also stained for the T cell activation marker CD69. Representative data for cells gated based on positive CD4 expression are shown. H. IL-2 concentration in supernatants of coculture experiments as described in Figure 4G. The IL-2 concentration was measured using a Bio-Plex assay. The data in B and D, and H are presented as the means ± SDs. Each dot represents one independent experiment. The data in F are representative of two independent experiments, and the data in G are representative of six independent experiments. The data presented in B, D, and H were analyzed by one-way ANOVA with Tukey’s multiple comparison test. n.s. = not significant, \**P* ≤ 0.05, \*\**P* ≤ 0.01, \*\*\**P* ≤ 0.001, \*\*\*\**P* ≤ 0.0001.

Taken together, these data support the existence of a cell-intrinsic heme-NRF2 signaling pathway that prevents the generation of functionally competent DCs.

### Phenotypical and functional DC depletion in the spleen of heme-exposed mice with genetic hemolytic anemia or Hmox1 deficiency

To investigate the potential involvement of heme-mediated effects on DCs in a pathophysiological context, we explored mouse models of diseases related to heme exposure. These investigations focused on the spleen, which has a dual function in mice during hemolysis, acting as a clearance organ for damaged RBCs and cell-free heme and as an extramedullary hematopoietic organ42,43. We thus expected that the spleen would mirror a heme-exposed BM-like microenvironment and replicate our cell culture model in vivo.

Intracellular heme levels are defined by the influx of extracellular heme into the cell (e.g., via erythrophagocytosis or the uptake of heme-protein complexes) and by intracellular heme degradation through heme oxygenases ^44^. The first model investigated in this study represents a condition of cell-intrinsic heme excess and is induced by conditional Hmox1 knockout. In these model mice, we deleted Hmox1 by tamoxifen treatment, which led to the progressive depletion of CD11c^+^/MHC class II^high^ DCs in the spleen. In contrast to the DCs, the pool of neutrophils was markedly enhanced nine weeks after tamoxifen treatment (**Figure 5A**), indicating that the observed depletion of DCs was not caused by nonspecific toxicity but rather by more specific hematopoietic deregulation.

**Figure 5:**
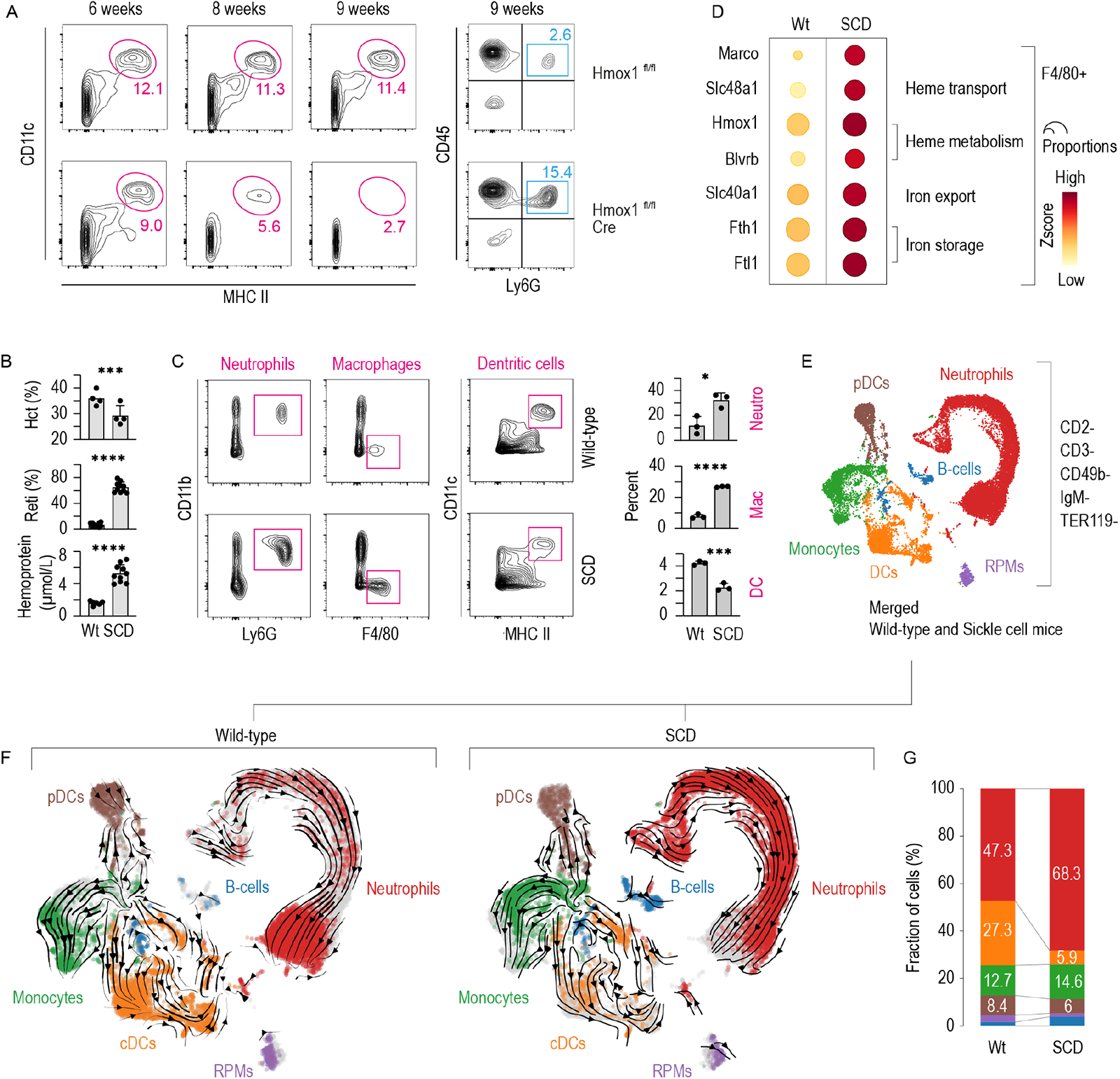
DC deficiency in the spleen of mice with Hmox1 deficiency or SCD. A. Left: Flow cytometry contour plots of spleen cell suspensions stained for CD11c and MHC class II from Hmox1^fl/fl^ Cre^neg^ and Hmox1^fl/fl^ Cre^pos^ mice at six, eight or nine weeks after the induction of target gene recombination by tamoxifen treatment. The cells were gated from live CD45^+^ cells. Right: Flow cytometry density plots of the spleen cell suspensions from the mice analyzed at nine weeks after tamoxifen induction. The suspensions were stained for CD45 and Ly6G. B. Hematocrit, reticulocytes, and plasma concentration of heme-adducted albumin in SCD mice and wild-type littermates. C. Left: Flow cytometry contour plots of spleen cell suspensions from SCD and wild-type mice stained for neutrophils (CD11b^+^ Ly6G^+^), macrophages (F4/80^+^), and DCs (CD11c^+^ MHC class II^high^). The cells were run on a spectral flow cytometer and gated as live CD45^+^ CD3^_^ CD19^_^ cells. Right: Percentages of neutrophils, macrophages, and DCs in three SCD and control spleens. D. Expression heatmap of heme and iron metabolism genes macrophages enriched populations isolated from the spleen of SCD and wild-type mice using anti-F4/80 antibody coated magnetic dynabeads. The dot size indicates the fraction of cells expressing the genes, and the color indicates the normalized Z-score. E. UMAP plot of pooled scRNA-seq data from two sequencing experiments with spleen cell suspensions from SCD and wild-type mice. Before sequencing, the spleen cells were enriched for DCs by negative selection using a DC isolation kit. The colors indicate the marker gene-derived cell-type attribution delineated in Supplementary Figure 5. F. UMAP plot of the scRNA-seq dataset split by genotype and colored by cell type. RNA velocity vectors are superimposed onto the UMAP plots with arrows indicating the velocity direction and magnitude (arrow length). G. The stack bar charts illustrate each cell type as a proportion of the whole number of cells per sample. The data in B and C are presented as the means ± SDs. Each dot represents one mouse. t-test (B, C); n.s. = not significant, \**P* ≤ 0.05, \*\**P* ≤ 0.01, \*\*\**P* ≤ 0.001, \*\*\*\**P* ≤ 0.0001.

The second model used, which represents cell-extrinsic heme excess, is a mouse model of SCD. Berkeley SCD mice exhibited pronounced hemolysis with lower hematocrit and more reticulocytes than their control littermates (**Figure 5B**). Intravascular hemolysis in SCD causes considerable free heme exposure, which leads to the accumulation of heme-adducted albumin in the plasma (**Figure 5B**). In the spleen of SCD mice spectral flow cytometry revealed expanded populations of F4/80^+^ macrophages and CD11b^+^/Ly6G^+^ neutrophils, whereas the CD11c^+^/MHC class II^high^ DC population was reduced (**Figure 5C**). These data suggest that the SCD mice exhibited deregulated homeostasis of the main phagocyte populations in the spleen through heme-induced expansion of macrophages and neutrophils and depletion of DCs.

To more specifically characterize these changes in the SCD mouse spleen, we performed scRNA-seq experiments with spleen cell suspensions from wild type and SCD mice that were (1) enriched for macrophages using anti-F4/80 antibody-coated magnetic beads or (2) enriched for DCs using a DC isolation kit based on the depletion of unwanted cells (mainly B cells and erythropoietic cells). Details of the F4/80^+^ macrophage scRNA-seq analysis of 21,170 cells are provided in **Supplementary Figures 3** and **4**. As expected from the results of the flow cytometry experiment (**Figure 5C**), red pulp and erythroblastic island macrophages were expanded in the spleen of the SCD mouse (**Supplementary Figures 3** and **4**). Consistent with an adaptive macrophage response, we found strongly increased expression of genes related to heme and iron metabolism and enhanced expression of the erythrophagocytic marker MARCO ^25^ in the hemolytic SCD mouse spleen (**Figure 5D**).

In the scRNA-seq analysis of enriched DCs, we used CellRank ^37^ to assign 9,419 wild-type and 10,660 SCD splenic cells to one of 10 macrostates (**Supplementary Figure 5A**). Based on a set of marker genes, every macrostate was assigned to one of six hematopoietic cell types (**Figure 5E** and **Supplementary Figure 5B/5C**). **Figure 5F** shows UMAP plots separated by genotype with arrows showing the RNA velocity vectors, which represent the potential differentiation trajectories. A significantly lower proportion of conventional DCs (cDCs) was found in the spleen of the SCD mouse than in the wild-type mouse (5.9% vs. 27.3%). Consistent with neutrophilia in the peripheral blood, we found a strongly left-shifted population of neutrophils in the SCD mouse spleen. This is visualized by the scattered distribution of the cells along the RNA velocity vector with more cells positioned close to the vetor origins indicating less mature cells (**Figure 5F**). Because categorial cell-type attribution based on marker genes as described above can be ambiguous and prone to bias, we analyzed splenic cells using a second method based on the use of RNA velocity data to assign each cell *n* probabilities of reaching a predefined number of *n*-terminal differentiation states (i.e., macrostates) ^37, 38^. We set the parameters to six terminal differentiation states, which reflected neutrophils, monocytes, plasmacytoid DCs (pDCs), cDCs, B cells, and red pulp macrophages. **Supplementary Figure 5D** shows the calculated probability for every cell in the data set to reach any of the six macrostates. **Supplementary Figure 5E** shows the cumulative absorbance probabilities split by genotype confirming that the SCD mouse spleen had a contracted cDC compartment.

To further define the subtype of DCs depleted from the SCD mice, we selected all DCs from the scRNA-seq dataset described in Figure 5 and regrouped them through another round of unsupervised Leiden clustering, which showed five distinct clusters (**Figure 6A**). We matched the gene expression profiles of each cluster with subtype-discriminating DC markers that were defined using the population comparison browser from ImmGen ^45^, which allowed us to attribute clusters 2, 0, and 1 to cDC1s, cDC2s, and pDCs respectively (**Figure 6B**). Additionally, we identified cluster 3 as a population of DCs highly similar to our in vitro-generated DCs in GM-CSF BM cultures, which showed notably strong expression of Ccr7 and Cd83 (**Figure 6C**). **Figure 6C** shows gene expression intensity UMAP projections for canonical DC genes that were not used for our Immgen based classification ^26, 36, 46–50^, demonstrating cDC1s with high expression of Xcr1, cDC2s expressing Itgam and Sirpa, and pDCs expressing Siglech. The data indicated that the observed depletion of DCs in the SCD mouse model could be attributed to a deficiency in cDC2s and the strong depletion of Ccr7^+^ Cd83^+^ mature DCs. We validated the scRNA-seq data across multiple mice by qRT-PCR analysis of independently enriched DCs from eight control and eight SCD animals, which revealed profound suppression of the signals for Cd74, H2-Ab1, and the DC maturation markers Cd83, Ccr7, and Cd40 in the SCD mouse spleens compared with the spleens of control littermates. The different genotype resulted in a clear segregation of wild-type and SCD mice in the hierarchical clustering analysis shown in **Figure 6D**. Collectively, these phenotype data suggest that SCD mice have severe deficiency of cDC2 and Ccr7^+^ Cd83^+^ mature DCs in the spleen.

**Figure 6:**
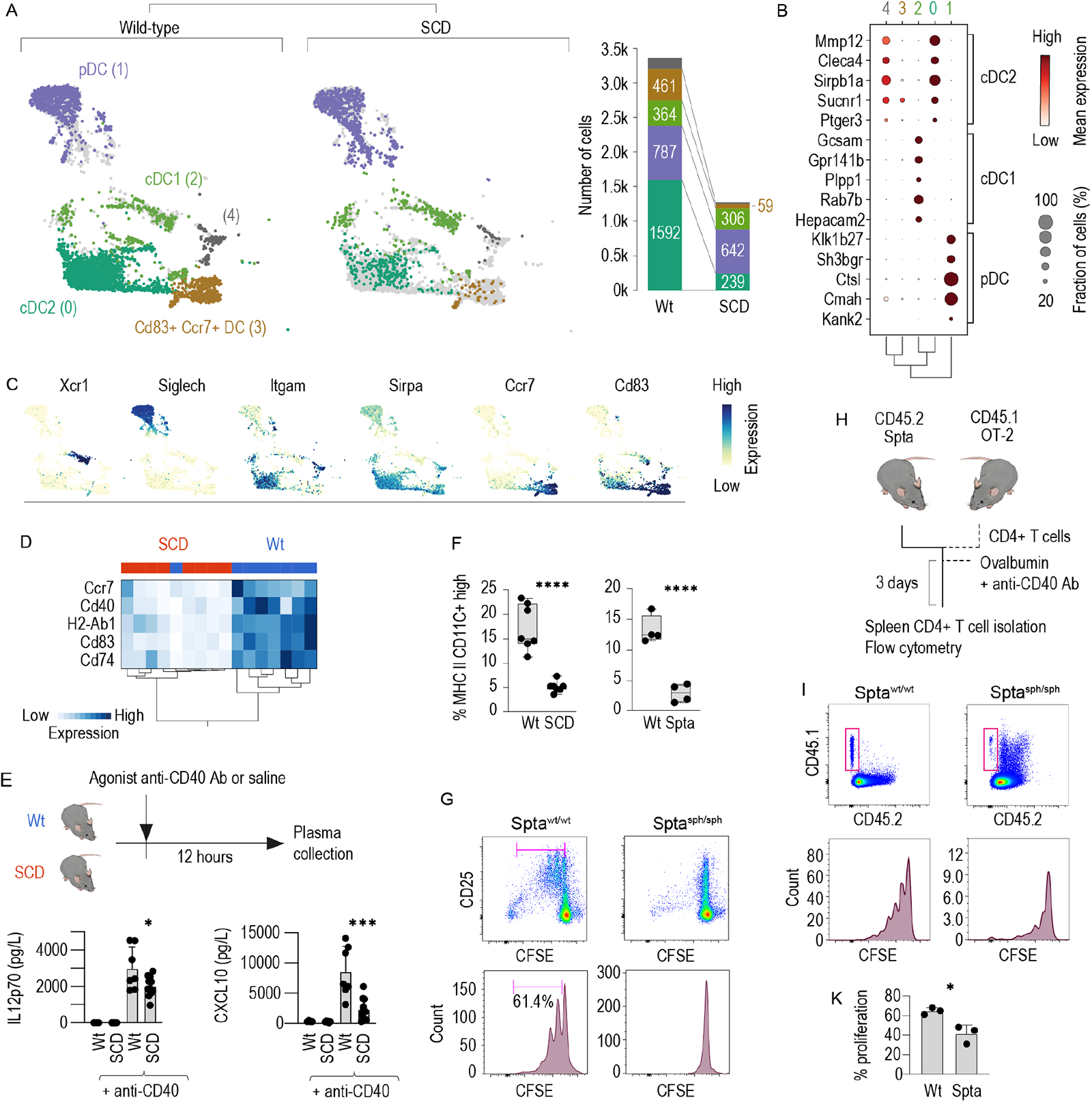
Functional DC depletion in the SCD spleen is caused by a loss of cDC2s and Ccr7^+^ Cd83^+^ mature DCs. A. Left: All DCs from the scRNA-seq dataset in Figure 5E/F were extracted and clustered using an unsupervised Leiden algorithm (numbers in parenthesis are cluster identities). For illustration of the differences, the dataset was separated by genotype (wild-type versus SCD). Right: The stack bar charts represent the number of cells per stratified for genotype. Numbers in parenthesis are cell counts. B. Cell-type attribution. The dot plot shows the normalized expression intensity (color) and the fraction of positive cells (size) for the top five discriminating ImmGen marker genes for cDC1s, cDC2s, and pDCs in each cluster. The dot size indicates the fraction of cells, the code color the mean expression. The dendrogram shows the hierarchical clustering based on the PCA components between the five clusters. C. UMAP intensity projections illustrate the expression of Xcr1, Siglech, Itgam, Sirpa, Ccr7, and Cd83 across the whole dataset of wild-type and SCD DCs. D. Hierarchical clustering analysis of the normalized mRNA expression intensities for Cd74, H2-Ab1, Ccr7, Cd40, and Cd83 in isolated splenic DC suspensions. Isolated DC samples were collected from the spleen of eight SCD mice and eight wild-type mice using a DC isolation kit and analyzed by qRT-PCR. Each column indicates one mouse. The almost perfect clustering within genotypes and the long dendrogram distance suggest highly different expressions across the set of marker genes in wild-type versus SCD mice. E. Plasma concentrations of CXCL10 and IL12p70 in SCD mice and wild-type littermates treated with saline (control) or an agonistic anti-CD40 antibody measured with a Bio-Plex assay. F. Percentage of live CD45^+^ cells splenic CD11c^+^ MHC class II^high^ DCs in SCD mice, Spta^sph/sph^ mice and their littermates measured by flow cytometry in DC enriched splenic cell suspension. G. Enriched DC suspensions from the spleen of Spta^sph/sph^ mice or wild-type littermates were pulsed with OVA_323-339_ (10 µg/ml) and cocultured with CFSE-labeled naïve CD4^+^ T cells isolated from OT-2 mice. The proliferation of CD4^+^ T cells was assessed with flow cytometry by evaluating the degree of CFSE dilution after four days in coculture in a 1/3 enriched DCs to T cells ratio. The cells were also stained for the T cell activation marker CD25. Representative data for cells gated based on positive CD4 expression are shown. H. Schematic representation of the adoptive T cell transfer experiments. CD4^+^ T cells were isolated from the spleen of CD45.1 x OT-2 mice, labeled with CFSE and injected i.v. into Spta^sph/sph^ mice or wild-type littermates. Subsequently, the mice were intravenously immunized with ovalbumin protein (75 µg) mixed with an agonistic anti-CD40 antibody. Three days later, the spleens were harvested, CD4^+^ T cells were negatively enriched and proliferation of CD4^+^ T cells was assessed by evaluating the degree of CFSE dilution. I. Top row: representative density plots of the CD45.1 and CD45. 2 expression in the enriched CD4 fraction isolated from the spleen of immunized mice. Cells were gated based on positive CD4 expression. Bottom row: CFSE dilution in cells gated based on positive CD45.1 and CD4 expression. K. Cumulative T cell proliferation data from three independent experiments. The data in F, E, and K are presented as the means ± SDs, and each dot represents one mouse. The data in G are representative of three independent experiments. t-test (F, K); one-way ANOVA with Tukey’s multiple comparison test (E); n.s. = not significant, \**P* ≤ 0.05, \*\**P* ≤ 0.01, \*\*\**P* ≤ 0.001, \*\*\*\**P* ≤ 0.0001.

To determine whether the quantitative DC deficiency in SCD mice observed by flow cytometry and scRNA-seq reflects an impairment of DC function, we administered an agonistic anti-CD40 antibody, which provides a very potent activation signal to DCs ^51^, and measured the cytokine response in the blood. Consistent with the decreased expression of marker genes for mature CD40^+^ DCs, we detected significantly attenuated plasma levels of IL12p70 and CXCL10 in SCD mice compared with littermates (**Figure 6E**), indicating lower DC activity.

We then aimed to determine whether the depletion of cDCs in the spleens of hemolytic mice coincided with a defective CD4 T cell priming. Because Berkley SCD mice have a mixed genetic background, the validity of functional immunological assays performed with this strain might be limited. Therefore, we used a mouse strain with hereditary spherocytosis that exhibits severe intravascular hemolysis resulting from a mutation in the spectrin alpha (Spta) gene ^52^. As demonstrated in **Figure 6F** and **Supplementary Figure 6A**, Spta^sph/sph^ mice also exhibited a reduced number of CD11c^+^/MHC class II^high^ DCs, suggesting that DC depletion in the spleen is a generic sequela of hemolysis.

Furthermore, we measured the antigen-presentation capacity of OVA_323-339_-pulsed enriched splenic DCs from Spta^sph/sph^ and wild-type littermates in a coculture assay with CFSE-labeled naive CD4^+^ T cells from OT-2 mice. The proliferation of CFSE-labeled lymphocytes was markedly impaired after coculture with DCs enriched from the spleens of Spta^sph/sph^ mice compared with coculture with DCs from the spleens of wild-type mice (**Figure 6G** and **Supplementary Figure 6B**). We subsequently evaluated the ability of splenic DCs to initiate a T cell response in vivo. To this end, we transferred CFSE-labeled CD4^+^ T cells from CD45.1 x OT-2 congenic mice into Spta^sph/sph^ or wild-type littermates and immunized the recipients with OVA protein mixed with an agonistic anti-CD40 antibody as the adjuvant (**Figure 6H**). After three days, we found that the proliferation of CD45.1^+^ CD4^+^ T cells in the spleens of Spta^sph/sph^ mice was significantly decreased compared to that of wild-type animals (**Figure 6I** and **6K**).

Overall, the results demonstrate that genetic hemolytic anemia alters the homeostasis of DCs in the spleen, leading to the depletion of cDC2s and mature Ccr7^+^ Cd83^+^ DCs. This altered phenotype is accompanied by reductions in cytokine release and antigen-specific CD4^+^ T cell expansion in the spleen, providing a novel mechanistic underpinning of hyposplenism-related secondary immunodeficiency in hemolytic anemia.

## DISCUSSION

Macrophages are pivotal defenders against the consequences of hemolysis ^15, 19^. This protection relies on the high capacity of specialized macrophages in the spleen and liver to remove damaged RBCs and cell-free heme from the blood and on functionally linked iron recycling pathways that promote adaptation to anemia ^15, 16, 53^. In this study, we explored the hypothesis that hemolysis-enhanced heme stress promotes the generation of homeostatic macrophages at the expense of DCs, which emerge from the same BM progenitors in a potentially competitive way. We found that the activation of NRF2 by heme skewed the differentiation trajectories of BM cultures towards red pulp macrophage-like cells that strongly expressed genes with an antioxidant function and roles in heme and iron metabolism. This physiologically meaningful adaptation—designed to increase disease tolerance during hemolytic stress—deterred the generation of mature DCs and impaired the capacity to prime antigen-specific CD4 T cells. Translational models of genetic hemolytic anemia reproduced the outcome of heme-deregulated myelopoiesis. In models of SCD and spherocytosis, we found that the expansion of heme- and iron-metabolizing macrophages in the spleen was accompanied by a profound deficiency in phenotypically and functionally mature DCs and impaired T cell priming capacity in the spleen.

We obtained the key finding of our study that heme skews myeloid differentiation trajectories in growth-factor supplemented mouse BM cultures. The supplementation of BM cultures with GM-CSF constitutes the first in vitro technique established for the production of DC-like cells, and these cells serve as a key model for basic science and therapeutic innovation in immunology ^32^. More recently, the phenotypically and functionally very diverse cellular output of these cultures, consisting of neutrophils, macrophages, and DCs, has called into question the utility of this system for producing homogeneous cultures of mature DCs for research and therapeutic purposes ^29, 54–56^. However, for our studies, the pronounced cell-type heterogeneity of GM-CSF BM cultures provided a unique opportunity for investigating how heme exposure impacts differentiation trajectories across a broad spectrum of potential cell fates within a single-cell culture dish.

To avoid the potential ambiguities of flow cytometry-based cell-type delineation, we decided to utilize a focused scRNA-seq strategy to validate our hypothesis that heme-exposure redirects the differentiation of myeloid precursors towards antioxidant and iron-recycling macrophages at the expense of DC generation. We leveraged the scRNA-seq approach with unbiased cell-type attribution based on RNA velocity analysis, which accounts for the continuity of differentiation states across divergent differentiation trajectories ^37, 38^. In our GM-CSF BM cell culture model, this high-dimensional analysis confirmed three trajectories for the generation of DCs, macrophages, and neutrophils. Heme exposure fundamentally redirected the differentiation pathway of GM-CSF BM cultures by inhibiting the production of functionally competent DCs and favoring macrophages with a Spic^+^ Hmox1^+^ Slc40a1^+^ red pulp macrophage-like phenotype. Interestingly, we found that the heme-suppressible DC population generated in vitro shared prominent phenotypic characteristics, such as high Ccr7 and Cd83 expression, with the DC population that was distinctly depleted in the SCD mouse spleen. Although the gene expression profile of the GM-CSF-induced DCs produced in vitro did not precisely match the full gene expression profile defined for any of the canonical DC subtypes found in vivo, it was previously noted that DCs produced with this culture method share principal characteristics with Ccr7^high^ migratory DCs and cDC2s, notably their strong capacity for the antigen-specific priming of naïve CD4^+^ T cells ^29, 56^. In a broader perspective, our study not only defines a new mechanism by which heme-exposure distorts DCs and immune homeostasis but it also provides a unique high-dimensional analysis of a classical cell culture model, which was and still is very popular amongst immunologists. As such, our data may provide a resource supporting novel interpretations of past and future studies.

We provide substantial evidence showing that the suppression of functionally competent DCs in our BM culture model was mediated by NRF2. NRF2 is the master regulator of the cellular response to oxidative stress. The finding that NRF2 activation by heme or synthetic agonists exerts such a strong effect on cell fate decisions was, therefore, unexpected. Based on prior literature, two principal mechanisms could explain this observation. By supporting the expression of genes involved in heme and iron metabolism, heme-activated NRF2 in myeloid precursors could push probabilistic cell fate decisions toward red pulp macrophage-like phagocytes ^25^. In addition, NRF2 suppresses several inflammatory signaling pathways that cooperatively promote the generation of mature DCs, such as the NFkB ^57^, STAT-1 ^58^, and STING-related interferon signaling pathways ^59^. Through this suppressive activity, heme-activated NRF2 could block the formation of mature DCs at any stage between early progenitor commitment and final maturation.

The intracellular translocation of heme by erythrophagocytosis or the receptor-mediated uptake of heme-protein complexes can suppress proinflammatory transcriptional networks in macrophages ^25, 58^, disrupt actin cytoskeletal dynamics to cause dysfunctional chemotaxis and phagocytosis ^60^, inhibit proteasome function ^61^, and enhance immunosuppressive heme metabolites produced by Hmox1 ^62^. In light of these prior studies and our own observations, it is likely that heme is a multifactorial driver of hyposplenism-related secondary immunodeficiency and, collectively, these results may stimulate exploring heme-directed therapeutics to reconstitute immune defence in patients with hemolytic anemia.

In summary, our results reveal a role of free heme as a disruptor of immune homeostasis at the interface of innate and adaptive immunity and delineate a novel mechanism underlying hyposplenism.

## METHODS

### Mice

#### SCD mice

Berkeley SCD mice ^63^ with the genotype (Tg(Hu-miniLCR α1GγAγδβS) Hba^-/-^, Hbb^-/-^ and carrying the sickle transgene (HBA-HBBs)41Paz were purchased from Jackson Laboratories. (Tg(Hu-miniLCR α1GγAγδβS) Hba^-/-^, Hbb^+/-^ and hemizygous for Tg(HBA-HBBs)41Paz were used as control.

#### Spherocytosis mice

Interbreeding heterozygous Spta^sph^ mice (B6.C3-Spta1^sph^/BrkJ X WB.C3-Spta1^sph^/BrkJ, The Jackson Laboratory) yielded spherocytic and wild-type offspring. Homozygous mice on a pure C57BL/6J or WB background were not viable. In contrast, F1 mice bred from heterozygous Spta^sph/wt^ mice that were separately maintained on the C57BL/6J and WB backgrounds were viable and not compromised.

#### Conditional Hmox1 knockout mice

Hmox1^tm1.1Hes^ ^64^ mice were obtained from Dr. Esterbauer (University of Vienna, Austria) and crossed with B6.Cg(UBC-cre/ERT2)1Ejb/J mice; Hmox1 knockout was induced by oral gavage of tamoxifen.

#### Other mice

B6.Cg-Tg(TcraTcrb)425Cbn/J (OT-2), B6.SJL-Ptprca Pepcb/BoyJ (CD45.1) and C57BL/6-CD45.1/T x B6.129S7-Rag1<tm1Mom>/J (CD45.1/Rag1^-/-^) mice were obtained from the Swiss Immunological Mouse repository (SwImMR). CD45.1 OT-2 mice were obtained by crossing CD45.1 mice with OT-2 mice. C57BL/6J mice were obtained from Charles River Laboratories, and Nrf2^-/-^ and wild-type littermates were obtained from Professor Yuet Wai Kan (University of California, San Francisco). All colonies were housed and bred in the specific-pathogen-free animal facility at the Laboratory Animal Services Center (LASC) of the University of Zurich in individually ventilated cages. Mice were housed under a 12/12-h light/dark cycle in accordance with international guidelines.

Mice of both sexes that were 12-16 weeks old were used for all experiments. All experimental protocols were reviewed and approved by the Veterinary Office of the Canton of Zurich (ZH161 2020). All animals were maintained at the animal facility of the University of Zurich (LASC) and treated in accordance with guidelines provided by the Swiss Federal Veterinary Office.

### BM cell cultures

BM cells were isolated by flushing the femurs and tibias of 8- to 10-week-old mice, followed by straining the BM through a 70-μm filter. Lin^—^ progenitor cells were isolated using a Lin^+^ cell depletion kit (Lin: CD5, CD11b, CD45R (B220), Gr-1 (Ly-6G/C), 7-4, and Ter-119; Miltenyi Biotec) and 4 & 10^5^ cells pro well were plated in tissue culture-treated six-well plates (Nunc Multidishes with UpCell Surface, Thermo Fisher) in 4 ml of complete RPMI-1640 medium (10% fetal calf serum (FCS), 1% L-glutamine) supplemented with 1% penicillin/streptomycin (P/S) and 20 ng/ml recombinant mouse GM-CSF or 100 ng/ml recombinant mouse M-CSF (PeproTech). For the M-CSF-supplemented cultures, half of the medium was removed on day 3 and new medium supplemented with M-CSF (100 ng/ml) was added. For the GM-CSF-supplemented cultures, fresh medium containing GM-CSF (2x, 40 ng/ml) was added on day 2. On day 3, half of the medium was removed and new medium supplemented with GM-CSF (20 ng/ml) was added. Cultures were treated on day 3 as indicated in the results section with heme-albumin, albumin (as vehicle/control), 10 μM ML-314 (Tocris), or 15 μM RA-839 (Tocris). The BM cells were harvested for analysis on day 7 from the thermosensitive cell culture plates after cooling to room temperature for 30 minutes.

### Porphyrin preparation for cell culture

Hemin (heme-chloride) and SnMP (tin mesoporphyrin) were obtained from Frontier Scientific (Newark). The porphyrins were dissolved in 10 ml of NaOH (100 mM) at 37°C, and 10 ml of 20% human serum albumin (CSL Behring AG) was then added. After 1 h of incubation at 37°C, the pH of the solution was adjusted to pH 7.4 using ortho-phosphoric acid, and the final volume was adjusted to 25 ml with a saline solution. The porphyrin-albumin solutions were filter-sterilized (0.22 µm) and used immediately ^65^.

### In vivo treatments

Mice were treated intravenously with 100 µg of agonistic anti-CD40 antibody (InVivoPlus, clone FGK4.5/FGK45). Twelve hours after the anti-CD40 antibody injection, blood samples were collected by cardiac puncture for cytokine measurement (Bio-Rad, Bio-Plex).

### Tamoxifen treatment of mice

Tamoxifen was prepared by dissolving in corn oil for a final concentration of 80 mg/ml. Hmox1^tm1.1Hes^ and littermates were given 60 μl (4.8 mg) tamoxifen solution once a day for five days by oral gavage.

### Preparation of spleen cell suspensions and cell enrichment

Spleens were harvested and mechanically disrupted in PBS and passed through a 70-μm cell strainer. The cell suspensions were then centrifuged, incubated in RBC lysis buffer (BioLegend) for 2 min at 37°C and centrifuged once more to obtain spleen cell populations devoid of mature erythrocytes. DCs were negatively enriched from spleen single-cell suspensions using a Dynabeads DC enrichment kit (Invitrogen) according to the manufacturer’s instructions. Macrophages were positively enriched from spleen single-cell suspensions using anti-F4/80 antibody (rat, Bioscience, Clone T45-2342)-coated Dynabeads (anti-rat IgG, Invitrogen) according to the manufacturer’s instructions.

### CD4**^+^** T cell isolation and CFSE labeling

CD4^+^ T cells were negatively enriched from spleen single-cell suspensions using a MagniSort CD4 enrichment kit (Invitrogen) according to the manufacturer’s instructions. Isolated CD4^+^ T cells were labeled with CFSE (Thermo Fisher) at 37°C for 20 min, washed with PBS and counted before use. The final purity confirmed by flow cytometry to be > 95%.

### OT-2 assay

A total of 10^4^ M-CSF- or GM-CSF BM cells or DCs enriched from spleen cell suspensions were plated in 96-well round-bottom plates, pulsed or not with 1 or 10 μg/ml OVA peptide 323–339 (Sigma) for 30 min at 37°C, and washed three times with PBS. Subsequently, 5 × 10^4^ CFSE-labeled naïve CD4^+^ T cells isolated from spleens of OT-2 mice were added to the BM cells or enriched splenic DCs in complete RPMI-1640 medium and cocultured at 37°C. CFSE dilution was assessed by flow cytometry and IL-2 concentration was measured in the supernatant by Bio-Plex assay after three or four days as specified.

### Transfer of GM-CSF phagocytes and OT-2-specific CD4^+^ T cells into CD45.1 Rag1^-/-^ mice

GM-CSF BM cells that were exposed to heme-albumin or albumin (vehicle/control) as described in the BM-derived cell cultures section were incubated in complete RPMI-1640 medium with 10 μg/ml OVA_323-339_ for 30 min at 37°C. The cells were washed three times in PBS and injected intravenously via the tail vein into Rag1^-/-^ mice expressing the CD45.1 antigen 2 h after the intravenous transfer of 1.5 × 10^6^ CFSE-labeled CD4^+^T cells isolated from OT-2 mouse spleens. Three days later, the CFSE dilution of splenic CD45.2^+^ CD4^+^ T cells was evaluated by flow cytometry.

### Transfer of OT-2 CD4^+^T cells into Spta^sph/sph^ and Spta^wt/wt^ mice

First, 1.5 × 10^6^ CFSE-labeled CD4^+^ T cells isolated from OT-2 × CD45.1 mouse spleens were injected intravenously via the tail vein into Spta^sph/sph^ mice and wild-type littermates (F1 C57BL/6J × WB). Two hours later, the mice were challenged intravenously with 75 μg of OVA protein (InvivoGen) mixed with 75 μg of agonistic anti-CD40 antibodies. The mice were sacrificed three days later, CD4^+^ T cells were enriched from the spleen by negative selection and the CFSE dilution of CD45.1^+^ CD4^+^ T cells was assessed by flow cytometry.

### Bio-Plex cytokine assays

The concentrations of IL-2, IL12-p70, and CXCL10 in coculture supernatants or plasma were determined using Bio-Plex Cytokine Assays (Bio-Rad). The assays were performed with a Bio-Plex 200 system (Bio-Rad), and the results were analyzed using Bio-Plex Data Pro software (Bio-Rad).

### Flow cytometry

Cells were preincubated with a LIVE/DEAD Fixable Near-IR cell stain kit (Invitrogen) and with Mouse BD Fc Block™ (≤ 1 μg/million cells in 100 μl, BD Biosciences) at 4°C for 10 min. The following antibodies were purchased from BD Bioscience: anti-CD45 (clone 30-F11), anti-CD19 (clone 1D3), anti-CD3 (clone 17A2), anti-CD25 (clone 7D4), anti-CD11c (clone HL3), and anti-I-A/I-E (clone M5/114.15.2). The following antibodies were purchased from BioLegend: anti-CD45.1 (clone A20), anti-CD45.2 (clone 104), anti-CD4 (clone GK1.5), anti-CD11b (clone M1/70), anti-CD115 (AFS98), anti-CD11c (clone N418), F4/80 (clone BM8), anti-Ly-6G (clone 1A8), and anti-CD69 (clone H1.2F3). The following antibody was purchased from eBioscience: anti-CD4 (clone GK1.5). Corresponding isotype-matched irrelevant specificity controls were purchased from BD, BioLegend, and eBioscience. Reticulocytes were stained with thiazole-orange (Sigma-aldrich). Multiparameter analysis was performed with an LSRFortessa analyzer (BD Biosciences), a SP6800 Spectral Analyzer (Sony) or an Aurora 5L spectral flow cytometer (Cytek). The data were analyzed using FlowJo software (version 10.7.1) and FCS express 7 (De Novo software).

### Sequencing-based workflows and data analysis

#### scRNA-seq data acquisition

*Multiplexed cell culture experiments:* Approximately 2 million cells per experimental condition were stained with 1 μg of TotalSeq™ B0301-B0308 anti-mouse Hashtag antibodies (BioLegend) according to the manufacturer’s instructions and pooled together at equal cell numbers. The pooled multiplexed sample was then processed according to the 10x Genomics Chromium Single Cell 3’ v3.1 Reagent Kits with Feature Barcoding Technology for Cell Surface Protein instruction guide. *Spleen cells:* Enriched DC and macrophage populations from the spleen of SCD mice and control littermates were run as separate samples according to the 10x Genomics Chromium Single Cell 3’ v3.1 Reagent Kits. For spleen single cell experiments, the sample and controls were loaded on the same chip and underwent library construction in parallel.

For all experiments, the sample volume was adjusted to a target capture of 10,000 cells and loaded on the 10x Genomics chromium next-GEM chip G to generate gel-beads-in-emulsion (GEMs). The GEM solution was placed in a Applied Biosystems Veriti 96 well thermal cycler for reverse transcription as described by the 10x Genomics instruction guide (53:00 min at 53°C followed by 5:00 min at 85°C). The resulting barcoded cDNA was then cleaned using Dynabeads MyOne Silane and amplified for 11 cycles (as recommended by the 10x Genomics user guide for a target cell recovery of >6,000 cells). After amplification, for multiplexed experiments, cDNA generated from polyadenylated mRNA for the 3’ gene expression library was separated from DNA from the Cell Surface Protein Feature Barcode for the Cell Surface Protein library with Dynabeads MyOne Silane and SPRIselect reagents based on size. The quality and concentration of both cDNA and DNA were assessed using High-Sensitivity D5000 ScreenTape (Agilent). All samples presented product sizes with a narrow distribution centered around 2000 pb and yielded between 50 and 800 ng cDNA (manually selecting products between 100-250 and 5000-6000 bp). cDNA and DNA were then subjected to enzymatic fragmentation, end repair and A-tailing. Adaptors were ligated to the fragmented cDNA and DNA, and the sample index was added during sample index PCR (set for 12 cycles, as recommended by the 10x Genomics user guide to correlate with a cDNA/DNA input of 12-150 ng). Library quality and concentration were assessed using High-Sensitivity D5000 ScreenTape (Agilent). All libraries showed an average fragment size of around 400 pb. For multiplexed runs, 3ʹ Gene Expression and Cell Surface Protein libraries were pooled at a ratio of 4:1 and sequenced using the Illumina NovaSeq 6000 system with a sequencing depth of 50,000 and 12,500 reads per cell, respectively, following the recommendations of 10X Genomics (paired-end reads, single indexing, read 1 = 28 cycles, i7 = 8 cycles, i5 = 0 cycles and read 2 = 91 cycles). For non multiplexed runs, 3ʹ Gene Expression libraries for each sample were pooled at an equimolar amount and sequenced using the Illumina NovaSeq 6000 system with a sequencing depth of 50,000 reads per cell, following the same recommendations of 10x Genomics.

#### scRNA-seq data analysis

The Cell Ranger Single-Cell Software Suite (version 4.0.0) was used for cDNA oligopeptide alignment, barcode assignment and UMI counting from fastq data from the Illumina sequencing. For each sample, the cell-containing droplets were filtered from the empty droplets, and this step was followed by the generation of an expression matrix using Cell Ranger Count (version 4.0.0). Demultiplexing of the cells within each sample was performed with the filtered matrix produced by Cell Ranger in R (version 4.4) using Seurat (version 3.2.3) and the HTODemux function (positive quantile set at 0.99). The resulting gene expression matrices were further analyzed with Python (version 3.8.8) using the Scanpy (version 1.6.0) ^66^, scVelo (version 0.2.2) ^38^ and CellRank (version 1.1.0) ^37^ libraries. Cells with a total number of expressed genes > 5,000 or < 500, a proportion of mitochondrial genes < 15%, or a proportion of ribosomal genes < 30% and genes expressed in < 20 cells were excluded from downstream analyses. After filtering, normalization was performed using the scran normalization algorithm implemented in the scran R package (version 1.18.3), which was followed by log normalization. For dimensional reduction, principal component analysis (PCA) of each sample was performed using the tl.pca function with the default settings from the Scanpy library. For cell culture samples, a shared nearest neighbor graph was built using the pp.neighbors function based on the first 15 PCAs or first 30 PCAs, respectively. Using the Leiden algorithm ^67^, cells were clustered and visualized in 2D using UMAP. For expression intensity projections, the cells were colored according to their log-normalized gene expression. Differentially expressed genes (p-value < 0.05 and a log2-fold change > 0.5) were determined using the tl.rank_genes_groups function from the Scanpy library with a pairwise Wilcoxon rank-sum test. For RNA velocity analysis, an expression matrix with spliced and unspliced transcript reads was calculated using the velocyto pipeline ^68^ by applying the Python script velocyto.py (version 0.17.17) to the Cell Ranger output folder. Using the scVelo library, RNA velocities were calculated by stochastically modeling the transcriptional dynamics of splicing kinetics, and the velocities were then projected as streamlines onto the low-dimensional UMAP embedding. With CellRank, cells were assigned to a predefined number of macrostates combining RNA velocity and transcriptional similarity information.

#### Pathway and transcription factor enrichment analysis of scRNA-seq data

GSEA was performed using the GSEApy library ^69^. With the enrichR function, which is based on EnrichR ^39^, we tested the enrichment for transcription factor terms using the TRRUST gene-set library ^40^. As the input, we used a gene list consisting of differentially expressed genes between the different conditions. Using gene lists ranked according to a combined score calculated with the tl.rank_genes_groups function from Scanpy, we performed pre ranked GSEA using the hallmark gene sets from the Molecular Signature Database (MSigDB). ^70^ For visualization, the terms were ordered according to their normalized enrichment score, and the running enrichment score for each term was plotted along the ranked genes as a heatmap.

Nrf2 target gene expression scores were calculated for each cell using the average expression of the Nrf2 gene set from the TRRUST gene-set library subtracted with the average expression of a randomly sampled reference set of similar length.

#### RT-qPCR analysis

Total RNA was isolated from spleen-cell suspensions using the RNeasy Mini Kit (Qiagen) according to the manufacturer’s instructions. Reverse transcription was performed with TaqMan reverse transcription reagents (Life Technologies). Real-time PCR was performed using Fast SYBR™ Green Master Mix (Applied Biosystems) to determine the expression levels of target genes using the primers listed in Table 1 shown below. Relative mRNA levels for experimental samples were calculated with 7500 Fast System Sequence Detection Software version 1.4 (Applied Biosystems) after normalization to the Hprt levels.

**Table 1:**
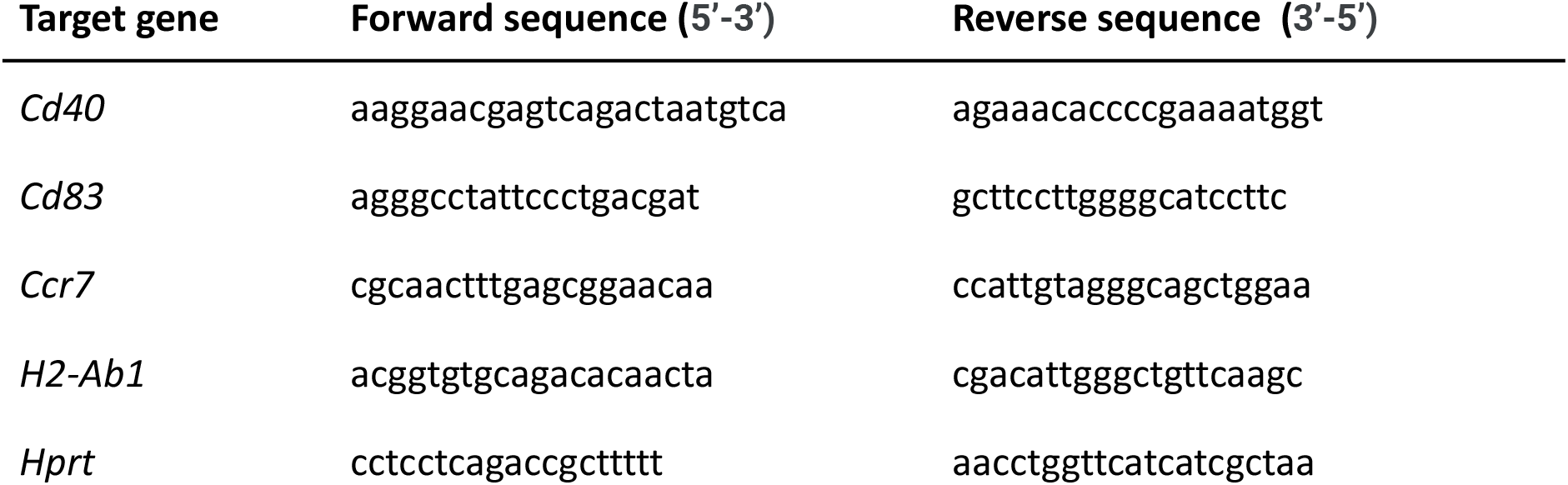
Sequences for PCR primers.

#### Free heme measurement

Mouse plasma samples were analyzed by SEC–high-performance liquid chromatography (SEC-HPLC) using an Agilent 1260 II HPLC attached to a quaternary pump and a photodiode array detector (DAD) (Agilent). Plasma samples and hemoglobin standards were separated on a Diol-300 (3 µm, 300 × 8.0 mm) column (YMC Co., Ltd.) with PBS (pH 7.4, Bichsel) as the mobile phase at a flow rate of 1 mL/min. For all samples, two wavelengths were recorded (λ = 280 nm and λ = 414 nm). From these data, we quantified peak areas of hemoglobin-haptoglobin complexes, free hemoglobin (dimers and tetramers), and other heme-adducted proteins ^71^.

#### Blood analysis

Hematocrit levels of blood samples of SCD and wild-type mice were measured by the Veterinary Laboratory of the University of Zurich.

#### Statistical analysis

Data plotting and statistical analysis were performed with Prism 9 (GraphPad), JMP 15 (SAS). For intergroup, we used a t-test or ANOVA with Tukey’s posttest as indicated in the figure legends. All data points are displayed in the graphs as mean ± standard deviation (n.s. = not significant, \**P* ≤ 0.05, \*\**P* ≤ 0.01, \*\*\**P* ≤ 0.001, \*\*\*\**P* ≤ 0.0001).

## AUTHOR CONTRIBUTIONS

FV designed the study, performed experiments, analyzed data, and wrote the paper. RMB implemented the RNA velocity analysis and analyzed data. MP performed the scRNA-seq experiments and analyzed data. KH performed experiments. AY performed the adoptive transfer experiments. AW and TG performed plasma analysis. RH wrote the paper. NS performed experiments. DJS designed the study, analyzed data, and wrote the paper.

## ACKNOWLEDGMENTS

This study was supported by the Vontobel Foundation (to FV), the Swiss National Science Foundation (310030_201202/1 to FV, MD-PhD scholarship to RMB, project 4221-06-2017, MD-PhD scholarship to MP, project 323530_183984, project grant 310030_197823 to DJS), and the Swiss Federal Commission for Technology and Innovation (project 19300.1 PFLS-LS to DJS).

## DATA SHARING STATEMENT

The datasets generated and/or analyzed during the current study are available from the corresponding author upon reasonable request.

