## Supplementary material for "Heme-stress activated NRF2 signaling skews fate trajectories of bone marrow cells from dendritic cells towards red pulp-like macrophages"

**SUPPLEMENTARY FIGURES AND RESOURCE TABLE**

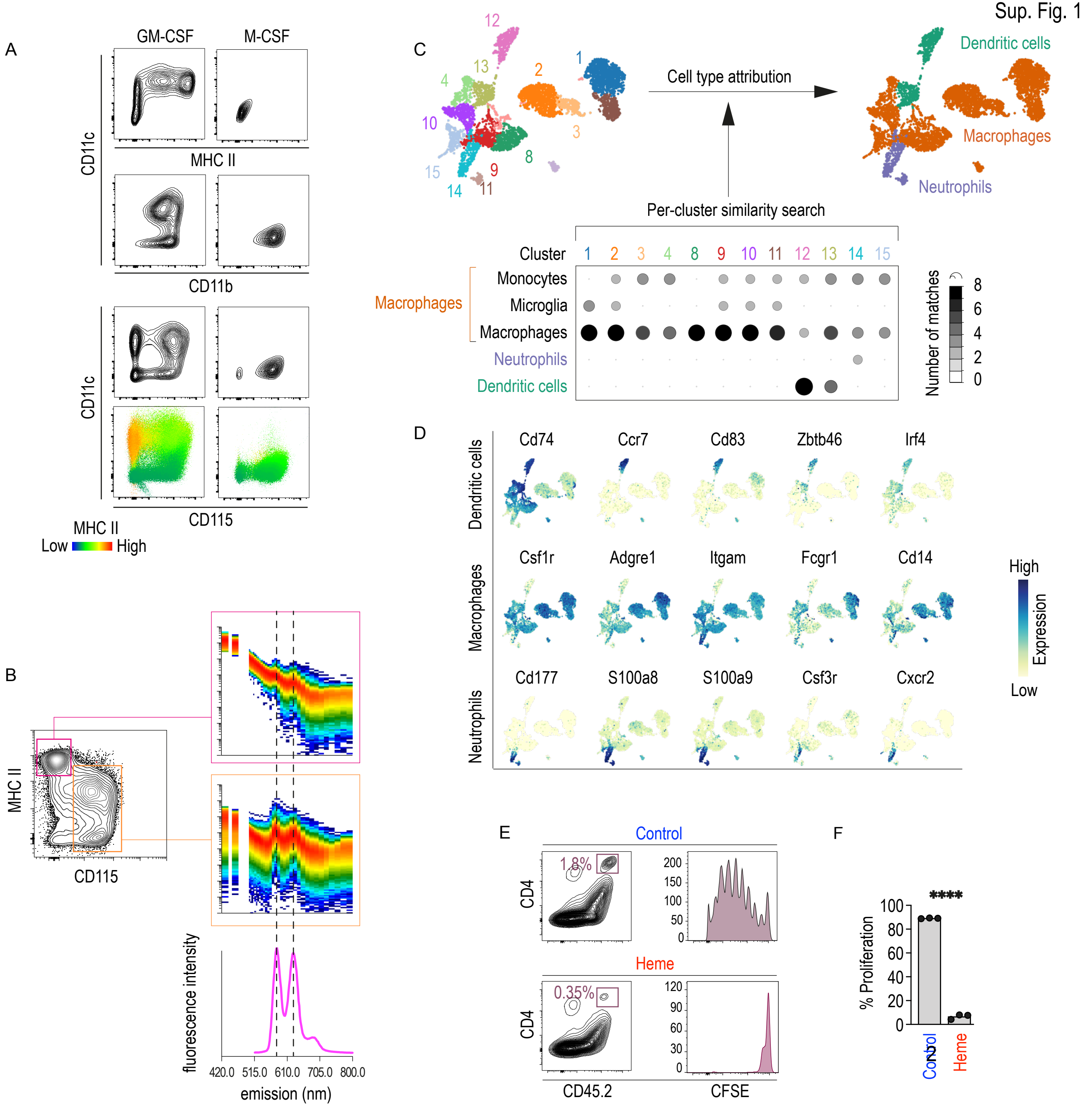
A
GM-CSF
M-CSF
CD11c
MHC II
CD11b
CD115
MHC II
Low
High
B
MHC II
CD115
fluorescence intensity
420.0 515.0 610.0 705.0 800.0
emission (nm)
C
Cell type attribution
Dendritic cells
Macrophages
Neutrophils
Per-cluster similarity search
Cluster
1 2 3 4 8 9 10 11 12 13 14 15
Monocytes
Microglia
Macrophages
Neutrophils
Dendritic cells
Number of matches
8 6 4 2 0
D
Dendritic cells
Cd74 Ccr7 Cd83 Zbtb46 Irf4
Macrophages
Csf1r Adgre1 Itgam Fcgr1 Cd14
Neutrophils
Cd177 S100a8 S100a9 Csf3r Cxcr2
High
Expression
Low
E
Control
CD4
1.8%
200 150 100 50 0
Heme
0.35%
120 90 60 30 0
CD45.2
CFSE
F
****
% Proliferation
100 80 60 40 20 0
Control
Heme

**Supplementary Figure 1**

A. Flow cytometry contour plots of GM-CSF- and M-CSF BM cultures. The cells were stained for CD11c, MHC class II, CD11b, and CD115.

B. SnMP fluorescence intensities were measured in GM-CSF BM cultures after 3h incubation with the SnMP porphyrin (100 µM)using a SP6800 Spectral Analyzer with the 640 nm laser inactivated.. The spectral plots show fluorescence emission intensities in macrophages defined as $CD115^{+}$ , MHC class $II^{low}$ (boxed in blue) and in DCs defined as $CD115^{-}$ MHC class $II^{high}$(boxed in magenta). The emission spectrum in macrophages matches the emission spectrum of SnMP measured in solution. Excitation was at 405 nm for flow cytometry and for the in solution fluorescence emission scan.

C. UMAP plot of the multiplexed scRNA-seq experiment described in Figure 1. Within the whole dataset, an unsupervised Leiden clustering identified 15 clusters. For cell-type attribution, we performed a per-cluster similarity search against > 15 million annotated single-cell transcriptomes from published studies using CellSearch (Bioturing). The dot plot shows the number of high-quality matches to published annotated cell populations for each cluster. The matching cell types were monocytes, microglia, macrophages, neutrophils, and DCs.

D. Expression intensity projections of canonical marker genes for DCs, macrophages, and neutrophils.

E. CFSE-labeled $CD4^{+}$ T cells isolated from the spleen of OT-2 mice and $OVA_{323\text{-}339}$ (10 µgr/ml)-pulsed GM-CSF BM cells that were exposed to heme or vehicle were injected intravenously via the tail vein into $Rag1^{-/-}$ mice expressing the CD45.1 antigen. Three days later, the spleens were harvested and the level of CFSE dilution was measured by flow cytometry. Left: representative contour plots showing the expression of CD4 and CD45.1. Right: CFSE dilution in cells gated based on $CD45.1^{+}$ $CD4^{+}$ T cells (red boxed).

F. Cumulative T cell proliferation data from three independent experiments.

The data in F are presented as the means ± SDs. Each dot represents one mouse. t-test; n.s. = not significant, *$P \leq 0.05$, **$P \leq 0.01$, ***$P \leq 0.001$, ****$P \leq 0.0001$.

Sup. Fig. 2

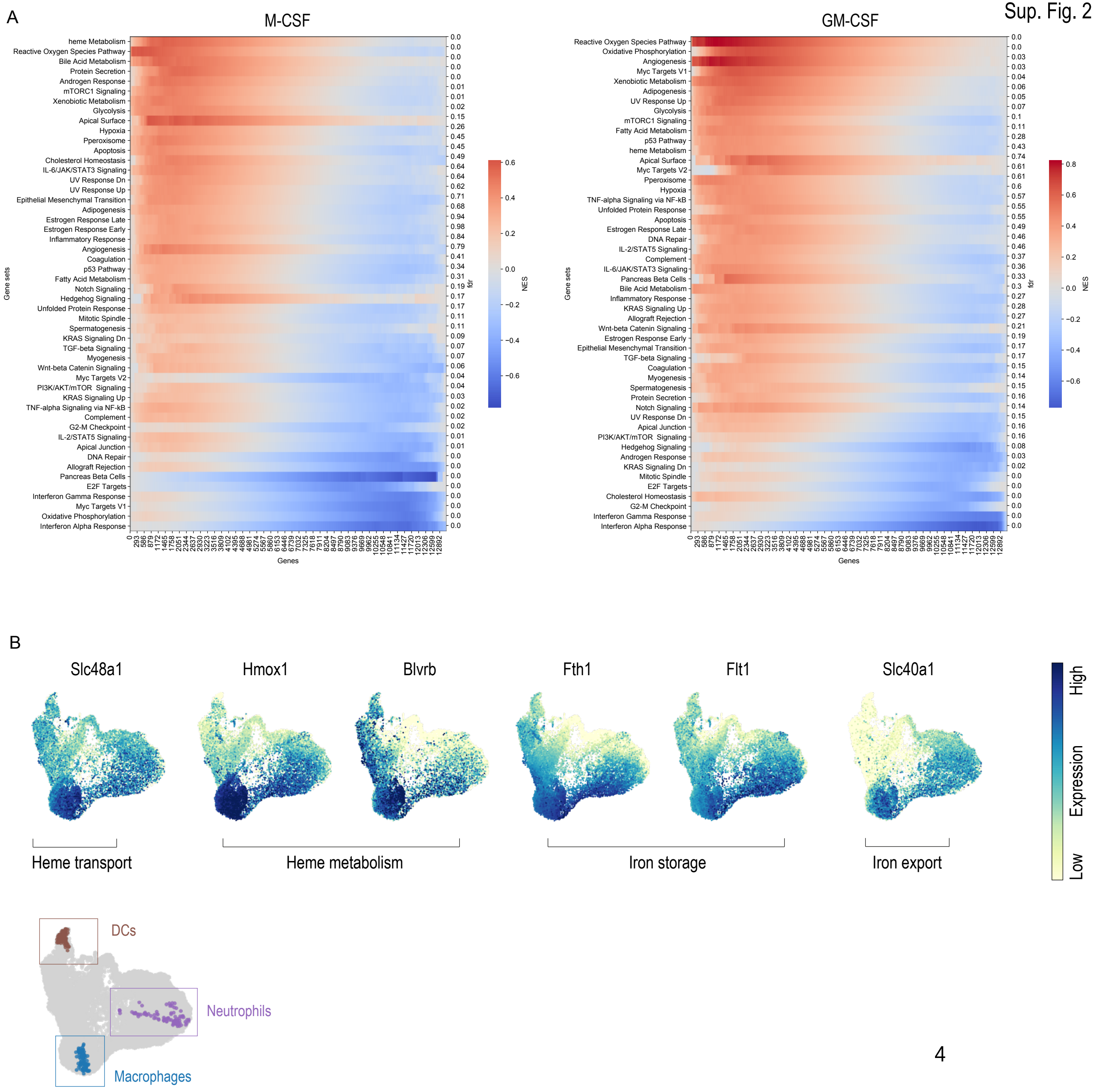
A
M-CSF
GM-CSF
heme Metabolism
Reactive Oxygen Species Pathway
Bile Acid Metabolism
Protein Secretion
Androgen Response
mTORC1 Signaling
Xenobiotic Metabolism
Glycolysis
Apical Surface
Hypoxia
Pperoxisome
Apoptosis
Cholesterol Homeostasis
IL-6/JAK/STAT3 Signaling
UV Response Dn
UV Response Up
Epithelial Mesenchymal Transition
Adipogenesis
Estrogen Response Late
Estrogen Response Early
Inflammatory Response
Angiogenesis
Coagulation
p53 Pathway
Fatty Acid Metabolism
Notch Signaling
Hedgehog Signaling
Unfolded Protein Response
Mitotic Spindle
Spermatogenesis
KRAS Signaling Dn
TGF-beta Signaling
Myogenesis
Wnt-beta Catenin Signaling
Myc Targets V2
PI3K/AKT/mTOR Signaling
KRAS Signaling Up
TNF-alpha Signaling via NF-kB
Complement
G2-M Checkpoint
IL-2/STAT5 Signaling
Apical Junction
DNA Repair
Allograft Rejection
Pancreas Beta Cells
E2F Targets
Interferon Gamma Response
Myc Targets V1
Oxidative Phosphorylation
Interferon Alpha Response
Gene sets
Genes
fdr
NES
Slc48a1
Hmox1
Blvrb
Fth1
Flt1
Slc40a1
Heme transport
Heme metabolism
Iron storage
Iron export
High
Expression
Low
B
DCs
Neutrophils
Macrophages

**Supplementary Figure 2**

A. GSEA of the differentially expressed genes in M-CSF or GM-CSF BM cultures (heme versus control) obtained by scRNA-seq analysis with annotation of all gene sets (rows) and the corresponding false discovery rate (referring to Figure 1). The heatmaps represent the magnitude of the running enrichment score per gene set category (red indicates positive enrichment, and blue indicates negative enrichment). The GSEA is based on the hallmark gene sets of the Molecular Signature Database (MSigDB).

B. Multiplexed scRNA-seq experiment of BM cells supplemented with GM-CSF as described in Figure 2. Top: the UMAP plots represent the expression intensity of genes associated with heme transport, heme metabolism, iron storage, and iron export. Bottom: UMAP plot showing the three terminal differentiated cell types as described in Figure 2B.

Sup. Fig. 3

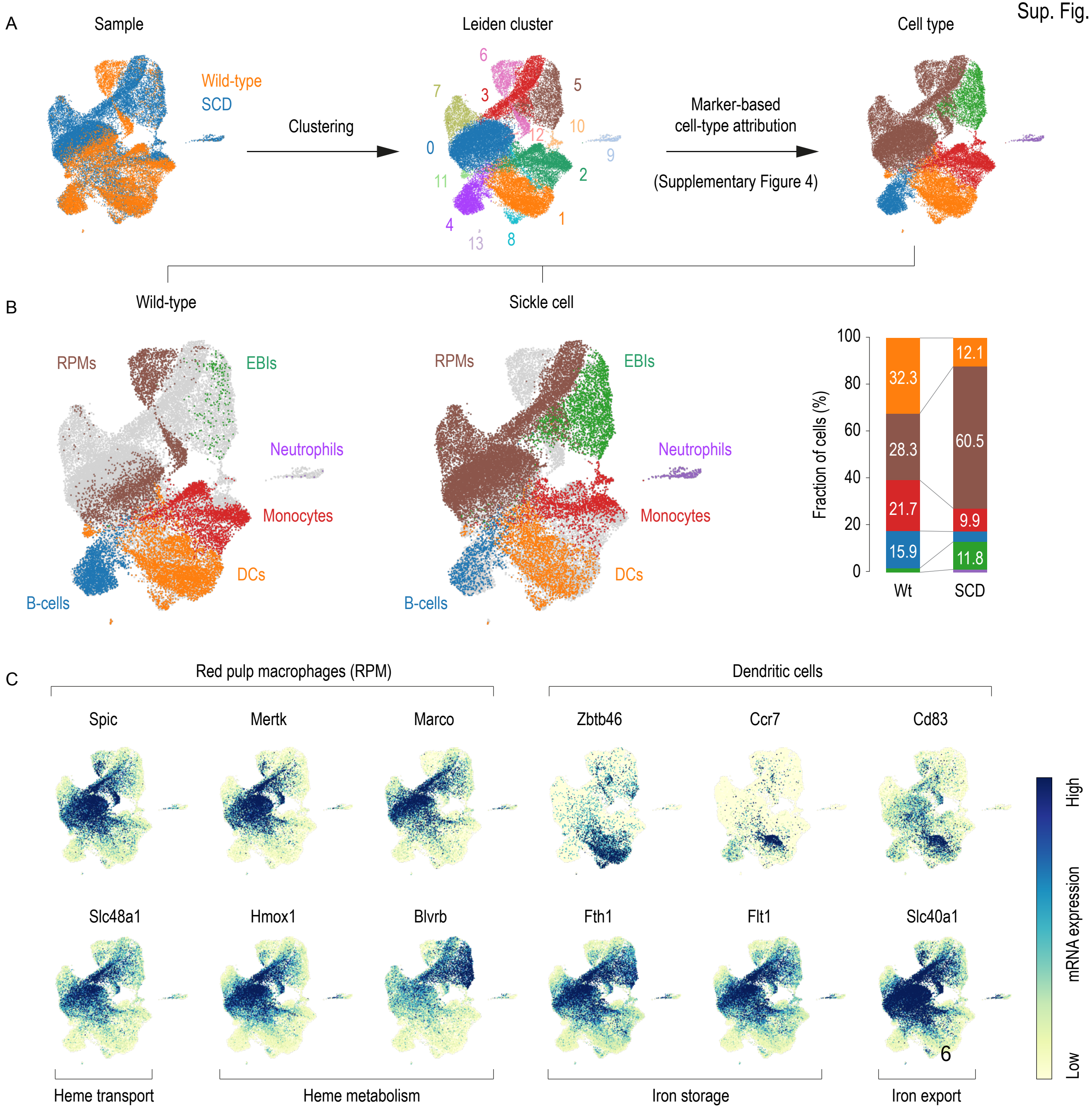
A
Sample
Wild-type
SCD
Clustering
Leiden cluster
0
1
2
3
4
5
6
7
8
9
10
11
12
13
Marker-based
cell-type attribution
(Supplementary Figure 4)
Cell type
B
Wild-type
Sickle cell
RPMs
EBIs
Neutrophils
Monocytes
DCs
B-cells
Fraction of cells (%)
100
80
60
40
20
0
32.3
28.3
21.7
15.9
12.1
60.5
9.9
11.8
Wt
SCD
C
Red pulp macrophages (RPM)
Dendritic cells
Spic
Mertk
Marco
Zbtb46
Ccr7
Cd83
Slc48a1
Hmox1
Blvrb
Fth1
Flt1
Slc40a1
High
mRNA expression
Low
Heme transport
Heme metabolism
Iron storage
Iron export

**Supplementary Figure 3**

A. UMAP plots of pooled scRNA-seq data from two sequencing experiments with spleen cell suspensions from an SCD mouse and a wild-type mouse. Before sequencing, the spleen cells were enriched for macrophages using anti-F4/80 antibody coated magnetic Dynabeads. For the whole dataset, unsupervised Leiden clustering identified 14 clusters that were attributed to six cell types (see Supplementary Figure 4).
B. Left: UMAP plots of the data split by strain (wild-type versus SCD). Right: The stack bar charts illustrate each cell type as a proportion of the whole number of cells per strain suggesting a massive expansion of red pulp macrophages (RPMs) and erythropoietic island macrophages (EBIs).
C. Top: Expression intensity projections of canonical marker genes for red pulp macrophages and DCs. Bottom: Expression intensity projections of genes associated with heme metabolism and iron metabolism.

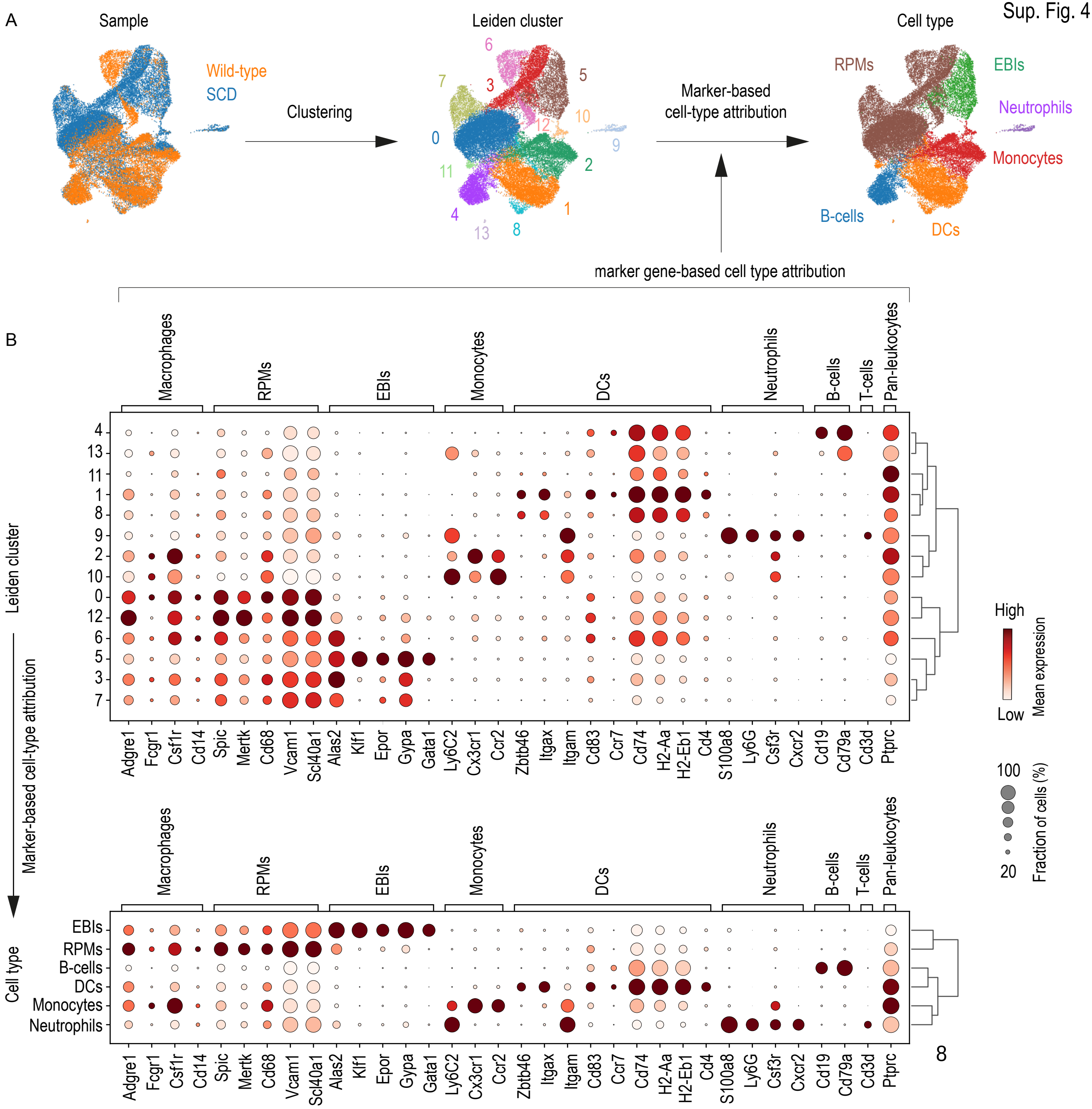
A
Sample
Wild-type
SCD
Clustering
Leiden cluster
Marker-based
cell-type attribution
Cell type
RPMs
EBIs
Neutrophils
Monocytes
B-cells
DCs
marker gene-based cell type attribution
B
Macrophages
RPMs
EBIs
Monocytes
DCs
Neutrophils
B-cells
T-cells
Pan-leukocytes
Leiden cluster
Marker-based cell-type attribution
Cell type
Adgre1
Fcgr1
Csf1r
Cd14
Spic
Mertk
Cd68
Vcam1
Scl40a1
Alas2
Klf1
Epor
Gypa
Gata1
Ly6C2
Cx3cr1
Ccr2
Zbtb46
Itgax
Itgam
Cd83
Ccr7
Cd74
H2-Aa
H2-Eb1
Cd4
S100a8
Ly6G
Csf3r
Cxcr2
Cd19
Cd79a
Cd3d
Ptprc
High
Low
Mean expression
100
20
Fraction of cells (%)

**Supplementary Figure 4**

Cell type attribution of the scRNA-seq experiment displayed in Supplementary Figure 3

A. UMAP plots of pooled scRNA-seq data from two sequencing experiments with spleen cell suspensions from an SCD mouse and a wild-type mouse. Before sequencing, spleen cells were enriched for macrophages using anti-F4/80 antibody coated magnetic Dynabeads. For the whole dataset, unsupervised Leiden clustering identified 14 clusters that were attributed to red pulp macrophages (RPM), erythroblastic island macrophages (EBIs), DCs, B lymphocytes, monocytes, and neutrophils.
B. Cell-type attribution. The dot plots show the normalized mean expression intensity (color) and the fraction of positive cells (size) for selected canonical marker genes in each Leiden cluster (top) and each identified cell type (bottom). The dendrograms show the hierarchical clustering based on the PCA components between the clusters (top) and the different cell types (bottom).

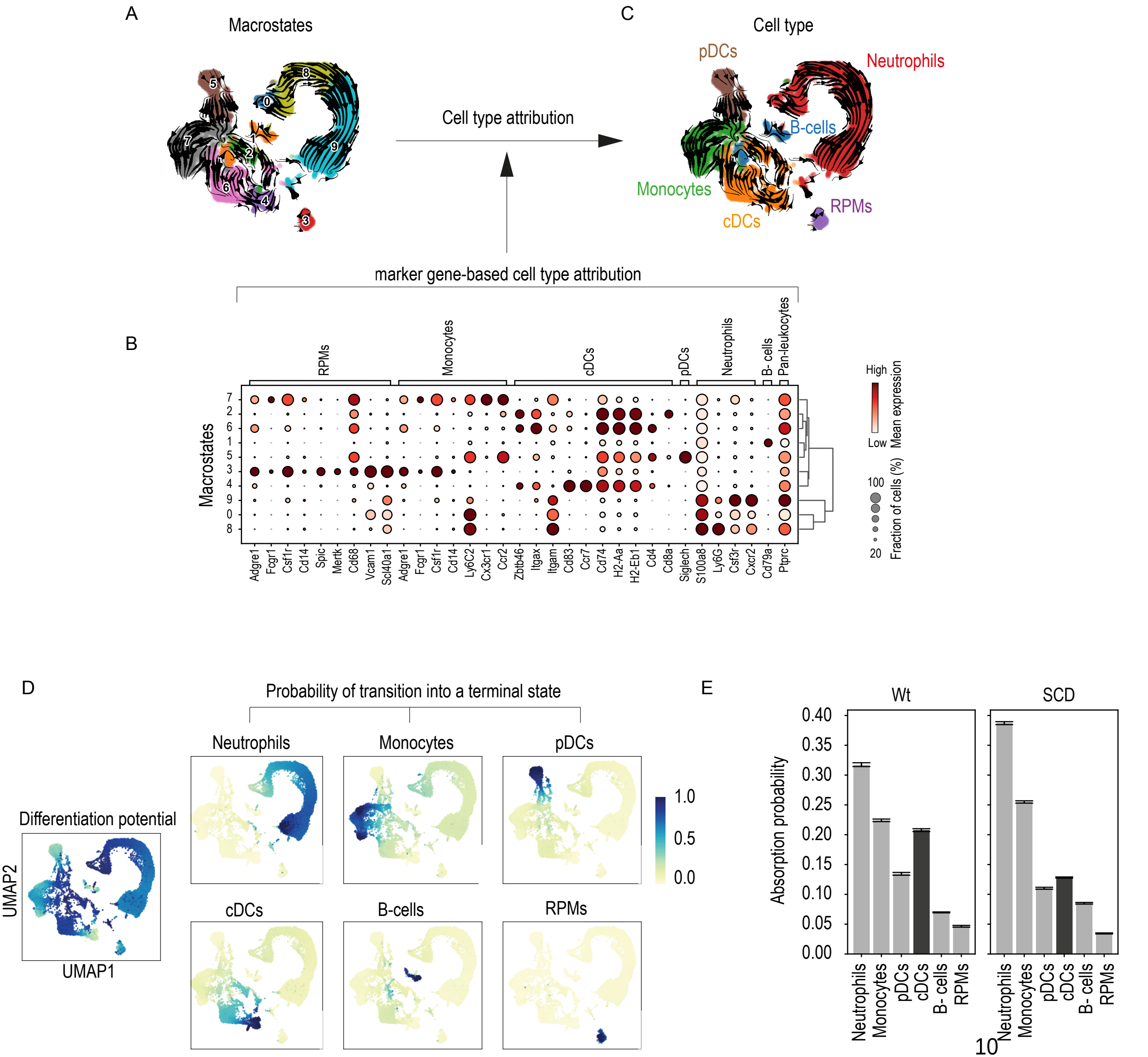
A
Macrostates
Cell type attribution
C
Cell type
pDCs
Neutrophils
B-cells
Monocytes
RPMs
cDCs
marker gene-based cell type attribution
B
RPMs
Monocytes
cDCs
pDCs
Neutrophils
B- cells
Pan-leukocytes
Macrostates
Adgre1 Fcgr1 Csf1r Cd14 Spic Mertk Cd68 Vcam1 Scl40a1 Adgre1 Fcgr1 Csf1r Cd14 Ly6C2 Cx3cr1 Ccr2 Zbtb46 Itgax Itgam Cd83 Ccr7 Cd74 H2-Aa H2-Eb1 Cd4 Cd8a Siglech S100a8 Ly6G Csf3r Cxcr2 Cd79a Ptprc
High
Low
Mean expression
100
20
Fraction of cells (%)
D
Probability of transition into a terminal state
Neutrophils
Monocytes
pDCs
cDCs
B-cells
RPMs
1.0
0.5
0.0
Differentiation potential
UMAP2
UMAP1
E
Wt
SCD
Absorption probability
0.40
0.35
0.30
0.25
0.20
0.15
0.10
0.05
0.00
Neutrophils
Monocytes
pDCs
cDCs
B- cells
RPMs

**Supplementary Figure 5**

A. scRNA-seq data of spleen cell suspensions from SCD and wild-type mice negatively enriched for DCs (see Figure 5). UMAP plots of the pooled scRNA-seq data (SCD and wild-type) colored by macrostates. The arrows indicate the calculated velocity.
B. Dot plot showing the mean expression levels of selected signature genes in each macrostate. The dot size indicates the fraction of cells expressing the genes, and the color indicates the per-gene normalized mean expression. The dendrogram represents the hierarchical clustering results based on the PCA components between the macrostates.
C. UMAP plots of the pooled scRNA-seq data (SCD and wild-type) colored by cell types.
D. The UMAP projections display the overall differentiation potential of every cell (left) and the transition probability of each cell into one of the six terminal states (right). For each terminal state a UMAP projection color coded representing the probability of each cell reaching this terminal state is shown.
E. Cumulative absorption probabilities across the whole experiment per cell type and genotype point to a reduced absorption probability for DCs in the SCD mouse. This plot aggregates and summarizes absorption probabilities from a single cell level for each experimental condition.

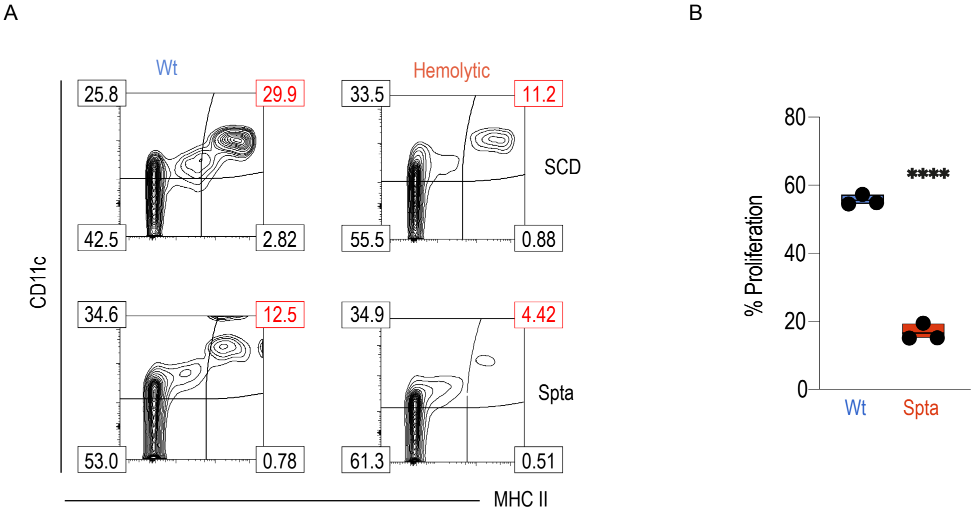
A
Wt
Hemolytic
25.8
29.9
33.5
11.2
SCD
42.5
2.82
55.5
0.88
CD11c
34.6
12.5
34.9
4.42
Spta
53.0
0.78
61.3
0.51
MHC II
B
% Proliferation
80
60
40
20
0
****
Wt
Spta

**Supplementary Figure 6**

A. Flow cytometry contour plots of splenic cell suspensions from SCD, $Spta^{sph/sph}$ mice, and their littermates. Representative cells were gated based on positive CD45 expression (cumulative data shown in Figure 6F). Percentage of DCs identified as CD11c and MHC class II positive cells are highlighted in red.
B. Cumulative T cell proliferation data from three independent experiments (corresponding to Figure 6G).

The data in B are presented as the means ± SDs. Each dot represents one independent experiment. t-test; n.s. = not significant, *$P \leq 0.05$, **$P \leq 0.01$, ***$P \leq 0.001$, ****$P \leq 0.0001$.

### Resource Table

#### FACS antibodies

| Reagent or Resource | Source | Identifier |
|---|---|---|
| Anti-CD115 BV605 | BioLegend | Cat # 135517 |
| Anti-CD115 PE-Cy7 | BioLegend | Cat # 135523 |
| Anti-CD11b APC-Cy7 | BioLegend | Cat # 101226 |
| CD11c APC | BioLegend | Cat # 117310 |
| CD11c PE | BD Pharmingen | Cat # 553802 |
| CD19 FITC | BD Pharmingen | Cat # 553785 |
| CD25 PE | BD Pharmingen | Cat # 558642 |
| CD3 BUV395 | BD Pharmingen | Cat # 740268 |
| CD4 PE | BioLegend | Cat # 100408 |
| CD4 eFluor 450 | ThermoFisher | Cat # 48-0041-82 |
| CD45 PE-Cy7 | BD Pharmingen | Cat # 552848 |
| CD45 Brilliant Violet 421 | BioLegend | Cat # 103134 |
| CD45.1 BV605 | BioLegend | Cat # 110739 |
| CD45.2 Pacific Blue | BioLegend | Cat # 109820 |
| CD69 APC | BioLegend | Cat # 104514 |
| F480 BV605 | BioLegend | Cat # 123133 |
| Ly6G PE | BioLegend | Cat # 127608 |
| Ly6G APC | BioLegend | Cat # 127614 |
| MHC2 BV421 | BD Pharmingen | Cat # 562564 |
| MHC2 Alexa Fluor 647 | BD Pharmingen | Cat # 562367 |
| TotalSeq™ B0301 Anti-mouse Hashtag antibodies | Biolegend | Cat # 155831 |
| TotalSeq™ B0302 Anti-mouse Hashtag antibodies | Biolegend | Cat # 155833 |
| TotalSeq™ B0303 Anti-mouse Hashtag antibodies | Biolegend | Cat # 155835 |
| TotalSeq™ B0304 Anti-mouse Hashtag antibodies | Biolegend | Cat # 155837 |
| TotalSeq™ B0305 Anti-mouse Hashtag antibodies | Biolegend | Cat # 155839 |
| TotalSeq™ B0306 Anti-mouse Hashtag antibodies | Biolegend | Cat # 155841 |
| TotalSeq™ B0307 Anti-mouse Hashtag antibodies | Biolegend | Cat # 155843 |
| TotalSeq™ B0308 Anti-mouse Hashtag antibodies | Biolegend | Cat # 155845 |
| Agonistic anti-CD40 antibody | InVivoPLus | Cat # BP0016-2 |
| Bio-Plex Pro™ Mouse Cytokine IP-10/ CXCL10 | Bio-rad | Cat # 12002244 |
| Mouse CD25/IL-2 R alpha DuoSet ELISA | R&D Systems | Cat # DY2438 |
| Bio-Plex Pro™ Mouse Cytokine IL12p70 | Bio-rad | Cat # 171G5011M |

#### Dyes

| Reagent or Resource | Source | Identifier |
|---|---|---|
| LIVE/DEAD™ Fixable Near-IR Dead Cell Stain Kit | Invitrogen | Cat # L10119 |
| Thiazole-orange | Sigma Aldrich | Cat # 390062 |

#### Chemicals, Peptides, and Recombinant proteins

| Reagent or Resource | Source | Identifier |
|---|---|---|
| EndoFit Ovalbumin (Chicken egg albumin; for in vivo use) | InvivoGen | Cat # 17E10-MM |
| Ovalbumin (323-339) (chicken, Japanese quail) | Sigma Aldrich | Cat # O1641 |
| RA-839 | Tocris | Cat # 5707 |
| ML-334 | Tocris | Cat # 5625 |
| Recombinant Murine GM-CSF | Peprotech | Cat # 315-03 |
| Recombinant Murine M-CSF | Peprotech | Cat # 315-02 |
| 20% Human Serum Albumin | CSL Berhing AG | Cat # 3665734 |
| Hemin | Frontier Scientific | Cat # H651-9 |
| SnMP (tin mesoporphyrin) | Frontiers Scientific | Cat # SnM321 |
| Phosphate buffered Saline (PBS) | Gibco | Cat # 10010-015 |

| | | |
|---|---|---|
| Penicillin-Streptomycin | Thermo Fisher | Cat # 15140-122 |
| RPMI Medium | Gibco | Cat # 11835-063 |
| Glutamax | Gibco | Cat # 35050-061 |
| MACS Buffer BSA Stock Solution | Miltenyi Biotec | Cat #130-091-376 |
| Dulbecco's MEM | Merck | Cat # 1469C |
| RBC Lysis Buffer (10X) | Biolegend | Cat #420301 |
| Fetales bovines Serum | Gibco | Cat # 10270-106 |
| Collagenase Type IV | Stemcell | Cat # 7427 |

#### Critical Commercial Assays

| Reagent or Resource | Source | Identifier |
|---|---|---|
| CellTrace™ Far Red Cell Proliferation Kit, for flow cytometry | ThermoFisher | Cat # C34564 |
| CellTrace™ Violet Cell Proliferation Kit, for flow cytometry | ThermoFisher | Cat # C34557 |
| UltraComp eBeads™ Compensation Beads | ThermoFisher | Cat # 01-2222-42 |
| Tru Stain FcxTmPLUS CD16/32, clone S17011E Isotype Rat IgG2b | Biolegend | Cat # 156604 |
| Dynabeads™ Mouse DC (Dendritic Cell) Enrichment | Invitrogen | Cat # 11429D |
| Dynabeads™ FlowComp™ Mouse CD4 Kit | Invitrogen | Cat # 11461D |
| MagniSort™ Mouse F4/80 Positive Selection Kit | Invitrogen | Cat # 8802-6863-74 |
| Lineage Cell Depletion Kit, mouse for 1×109 total cells | Miltenyi Biotec | Cat # 130-090-858 |

#### Experimental Models: Organisms/Strains

| Reagent or Resource | Source | Identifier |
|---|---|---|
| Mouse: C57BL/6NCrl | Charles River | 027C57BL/6 |
| Mouse: Hba<tm1Paz> Hbb<tm1Tow> Tg(HBA-HBBs)41Paz/J | The Jackson Laboratory | Jax # 003342 |
| Mouse: Spta1 B6.C3-Spta1<sph>/BrkJ | The Jackson Laboratory | Jax # 000450 |
| Mouse: WB.C3-Spta1 sph /BrkJ | The Jackson Laboratory | Jax # 000454 |
